# Temporal single-cell transcriptomes of zebrafish spinal cord pMN progenitors reveal distinct neuronal and glial progenitor populations

**DOI:** 10.1101/2021.04.28.441874

**Authors:** Kayt Scott, Rebecca O’Rourke, Caitlin C. Winkler, Christina A. Kearns, Bruce Appel

## Abstract

Ventral spinal cord progenitor cells, which express the basic helix loop helix transcription factor Olig2, sequentially produce motor neurons and oligodendrocyte precursor cells (OPCs). Following specification some OPCs differentiate as myelinating oligodendrocytes while others persist as OPCs. Though a considerable amount of work has described the molecular profiles that define motor neurons, OPCs, and oligodendrocytes, less is known about the progenitors that produce them. To identify the developmental origins and transcriptional profiles of motor neurons and OPCs, we performed single-cell RNA sequencing on isolated pMN cells from embryonic zebrafish trunk tissue at stages that encompassed motor neurogenesis, OPC specification, and initiation of oligodendrocyte differentiation. Downstream analyses revealed two distinct pMN progenitor populations: one that appears to produce neurons and one that appears to produce OPCs. This latter population, called Pre-OPCs, is marked by expression of *GS Homeobox 2* (*gsx2)*, a gene that encodes a homeobox transcription factor. Using fluorescent in situ hybridizations, we identified *gsx2*-expressing Pre-OPCs in the spinal cord prior to expression of canonical OPC marker genes. Our data therefore reveal heterogeneous gene expression profiles among pMN progenitors, supporting prior fate mapping evidence.

**Highlights:**

- Single-cell RNA sequencing reveals the developmental trajectories of neurons and glia that arise from spinal cord pMN progenitor cells in zebrafish embryos
- Transcriptionally distinct subpopulations of pMN progenitors are the apparent sources of neurons or oligodendrocytes, consistent with fate mapping data
- *gsx2* expression marks pMN progenitors that produce oligodendrocyte lineage cells

## Introduction

The vertebrate central nervous system (CNS) is composed of intricately organized arrays of distinct neurons and glia, which allow organisms to perceive and interact with their environment. Before these functional networks are established, neural progenitors are first spatially organized into distinct progenitor populations. This process, known as patterning, occurs as neural progenitors receive discrete positional cues created by combinatorial gradients of morphogens. For example, the ventral spinal cord is patterned by a ventral-to-dorsal gradient of the morphogen Shh (Briscoe and Ericson, 2001; Briscoe et al., 1999; Dessaud et al., 2008). Following CNS patterning, neural progenitors first produce neuronal cell types and then switch to producing glial cell types later in development (Kessaris et al., 2001; Miller and Gauthier, 2007). This change in fate is accompanied by alterations in intercellular and extracellular signaling that lead to the inhibition of neurogenesis and activation of gliogenesis (Kessaris et al., 2001; Miller and Gauthier, 2007).

An example of these events occurs in the ventral spinal cord, where high levels of Shh ligand secreted by cells in the notochord and floor plate are required to specify pMN progenitors, which express the basic helix loop helix transcription factor Olig2 (Dessaud et al., 2007; Dessaud et al., 2010; Echelard et al., 1993; Marti et al., 1995; Roelink et al., 1994). After specification pMN progenitors first produce motor neurons and then oligodendrocyte precursor cells (OPCs), some of which differentiate into myelinating oligodendrocytes (Lu et al., 2000; Noll and Miller, 1993; Novitch et al., 2001; Warf et al., 1991; Zhou and Anderson, 2002; Zhou et al., 2000). Notably, the switch from motor neuron to OPC specification requires a transient increase in ventral Shh signaling activity (Danesin and Soula, 2017). Though we understand many of the signaling pathways and transcription factors required for motor neuron and OPC specification, it is less clear how pMN progenitors are assigned to neuronal or glial fates.

Numerous lineage tracing studies have shown that most clonally related cells consist either of neurons or glia but not both (Luskin et al., 1988; Luskin et al., 1993; McCarthy et al., 2001), suggesting that neurons and glia are produced by different progenitors. Consistent with this interpretation, our fate mapping and time-lapse imaging studies in zebrafish revealed that motor neurons and OPCs arise from different pMN progenitor cells that initiate *olig2* expression at different times (Ravanelli and Appel, 2015). These results imply that different neural progenitors express genes that specify them for either neuronal or glial fates, but our knowledge of the gene functions that determine these distinct fates remains incomplete.

Here, we set out to identify transcriptional profiles of oligodendrocyte and motor neuron progenitors in the developing zebrafish spinal cord. We utilized single-cell RNA sequencing (scRNA-seq) of isolated pMN cells, which allowed us to characterize genes expressed by pMN cells at key developmental time points. These data revealed two distinct populations of pMN progenitors: one that expressed genes characteristic of neuronal fates and another, called Pre-OPCs, that appeared fated to give rise to OPCs. Fluorescent in situ RNA hybridization experiments confirmed the presence of neuronal progenitors and Pre-OPCs in the spinal cord prior to formation of OPCs. Our data indicate that pMN progenitors are allocated to distinct glial and neuronal lineages prior to the onset of OPC specification and reveal new genes that might be important for the earliest steps in formation of oligodendrocyte lineage cells.

## Results

### A developmental single-cell RNA expression atlas of zebrafish pMN progenitors and their derivative cell types

In zebrafish embryos, pMN progenitors produce the majority of spinal cord motor neurons by 25 hours post fertilization (hpf) (Myers et al., 1986) and most spinal cord oligodendrocyte precursor cells (OPCs) between approximately 28 and 42 hpf (Ravanelli and Appel, 2015) (Fig. 1A). Some OPCs then undergo differentiation into pre-oligodendrocytes (Pre-OL), denoted by expression of *myrf*, a transcript that encodes the transcription factor Myelin regulatory factor (Myrf). By 72 hpf a majority of the Pre-OLs mature into myelinating oligodendrocytes, marked by expression of *mbpa*, which encodes Myelin basic protein (MBP) (Brösamle and Halpern, 2002). To gain insight to the identity of neuronal and glial progenitors and gene expression changes that accompany formation of neurons and glia, we sorted *olig2:*EGFP^+^ cells obtained from 24, 36 and 48 hpf transgenic embryos to capture periods of neuron production, OPC production, and initiation of OPC differentiation, respectively (Fig. 1B). At each stage, we removed cranial tissue from our samples and combined the trunks and tails for subsequent cell dissociation, FACS and sequencing (Fig. 1B). Using unsupervised Seurat clustering we uncovered 28 distinct clusters and determined cell population identity within an integrated data set containing cells from all collected stages by assessing genetic markers within each cluster (Fig. 2; Fig. S1). Mapping the developmental timepoint origin of each cell within the integrated data set showed that some clusters included cells from only 24 hpf or 48 hpf embryos, indicative of unique progenitor populations present at 24 hpf and differentiated cell populations at 48 hpf (Fig. 2B). Most cells expressing markers characteristic of S, G2 and M phases of the cell cycle were isolated at 24 hpf and 36 hpf stages whereas most cells in G1 phase were from 48 hpf embryos (Fig. 2B,C). Together, these observations indicate that our experimental design captured cells representing a progression from neural progenitors to differentiated cells.

**Fig 1.**
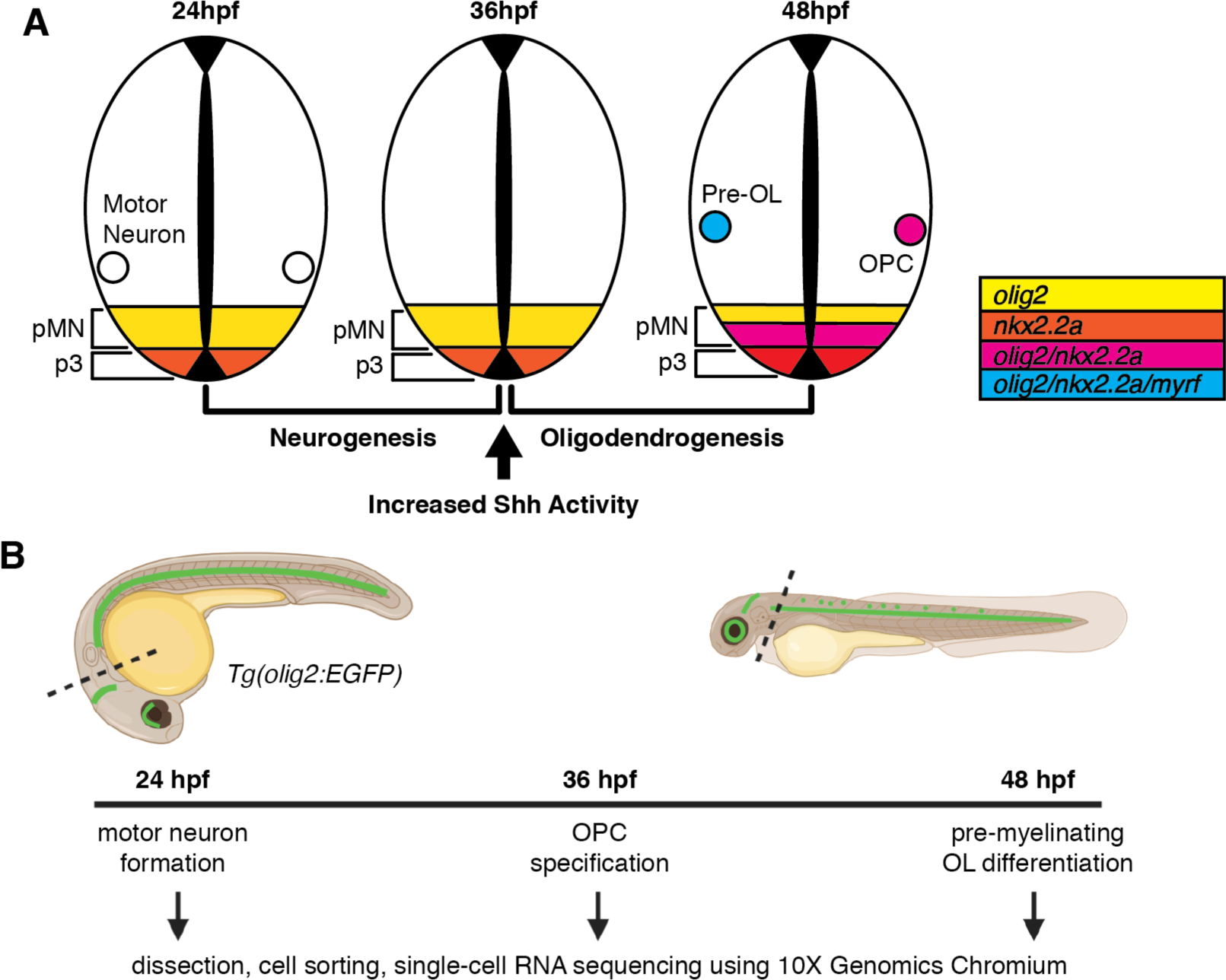
Zebrafish pMN cell development and cell isolation and scRNA-seq strategy. (A) Schematic of transverse sections of the zebrafish spinal cord depicting p3 and pMN ventral progenitor domains. The pMN domain is marked by expression of *olig2* and first produces motor neurons. Following a transient increase in Shh activity, *nkx2*.*2a* (red) is co-expressed (orange) with *olig2* by ventral pMN progenitors that subsequently develop as OPCs. (B) Schematic of the experimental design by which the posterior tissue of 24, 36 and 48 hpf *Tg(olig2:EGFP)* embryos were collected to isolate *olig2:*EGFP^+^ cells via FACS, followed by scRNA-seq using collected *olig2:*EGFP^+^ cells.

**Fig 2.**
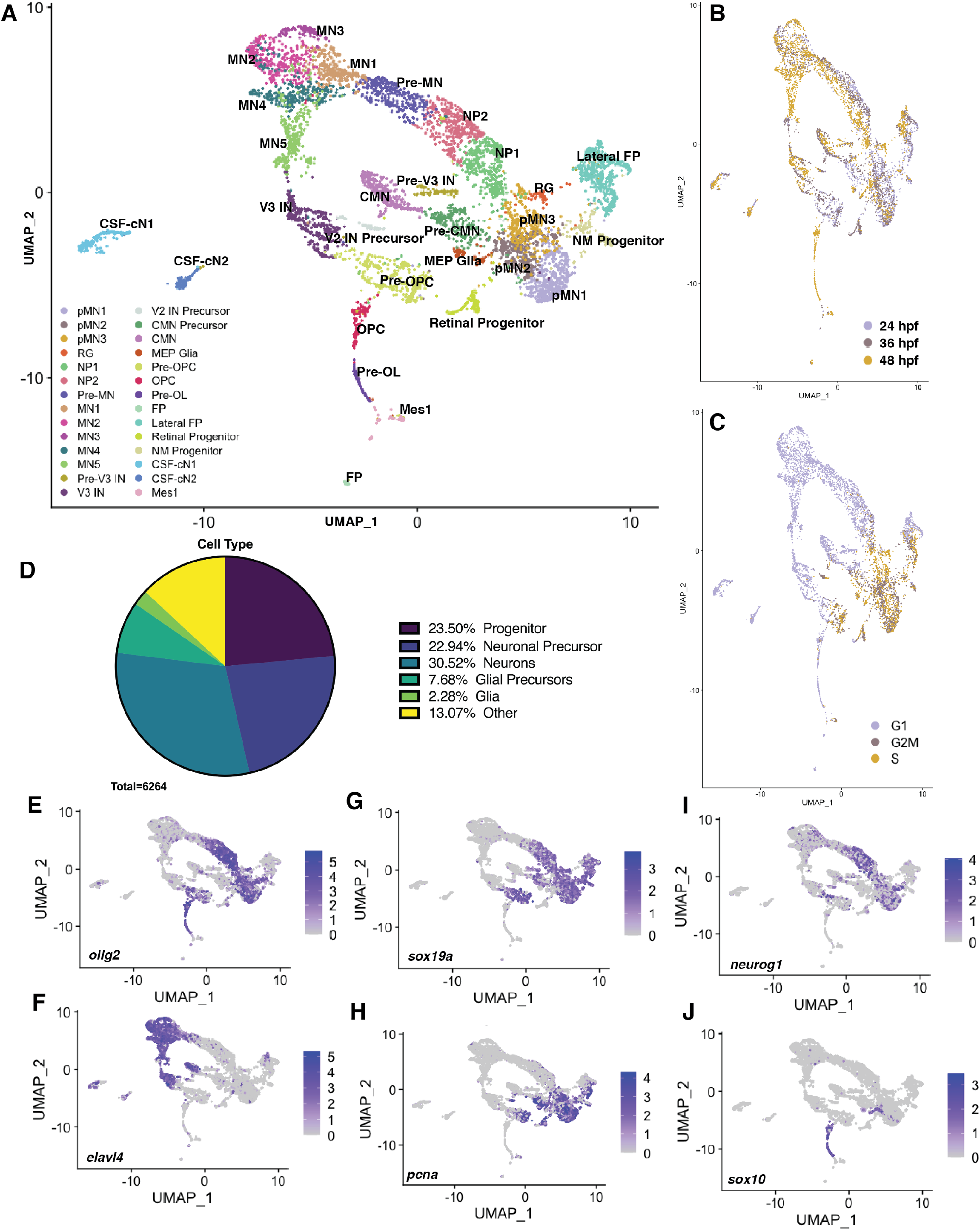
pMN cell diversity through embryonic zebrafish development. (A) Unsupervised UMAP of integrated scRNA-seq data from *olig2:*EGFP^+^ spinal cord cells obtained from 24, 36 and 48 hpf *Tg(olig2:EGFP)* embryos. Seurat analysis identifies 28 clusters composed either uniquely of or as a combination of cells from each collected time point. Each point represents one cell (n=6489). (B) Distribution of cells from each time point. (C) A majority of clusters contained cells in G1 phase and a few populations, likely representing progenitors, were in S and G2M phase. (D) Proportions of pMN gross cell type identities at all time points. A majority of isolated pMN cells are neurons or neuronal precursors; ∼24% are progenitors and ∼10% belong to glial lineages. (E-J) UMAP feature plots of selected transcripts. Cells are colored by expression level (gray is low, purple is high). o*lig2*^*–*^ *elav4*^*+*^ cells represent post-mitotic neuronal populations (E,F). Co-expression of *olig2, sox19a* and *pcna* designate pMN progenitors (G-H) and expression of *neurog1* in a subset of *olig2*^*+*^ *sox19*^*+*^ and/or *pcna*^*+*^ cells represent neuronal progenitors and precursors (I). *sox10* expression denotes glial cells (J).

Within the integrated dataset the majority of cells could be classified as neural progenitors, committed neuronal precursors, and neurons, with fewer cells belonging to glial lineages (Fig. 2D).

Notably, our dataset included numerous cell clusters that did not express *olig2* (Fig. 2E). Most of these *olig2*^*–*^ cells expressed *elavl4*, which marks post-mitotic neurons (Fig. 2F). We attribute the presence of *olig2*^*–*^ cells to the long perdurance of EGFP, which has a half-life of approximately 15 hours (Danhier et al., 2015). Thus, *olig2*^*–*^ cells likely represent cells that previously expressed *olig2* transcripts but then terminated transcription during neuronal differentiation. These data also included cells that we identified as cranial motor neuron (CMN) precursors, CMNs and Retinal Progenitors denoted by *phox2bb* expression (Pattyn et al., 2000), which likely were among the sorted cell populations due to imprecision in our cranial dissections (Fig. S1F). Within the original integrated data clustering, CMN precursors were grouped together with Motor Exit Point (MEP) glia (Fig. S2A,B). However, within this cluster there were distinct expression profiles that led us to sub-cluster this group, splitting it into CMN precursors marked by *sox19a* and *phox2bb* (Pattyn et al., 2000) and MEP glia, which expressed *sox10* and *foxd3* (Fig. S2C-F) (Fontenas and Kucenas, 2021; Smith et al., 2014). The additional sub-clusters within the CMN precursor group were based on differences in cell cycle and temporal origins (Fig. S2G-I). Of the cells that expressed *olig2*, a majority also expressed *sox19a* and *pcna*, indicative of cells with a neural progenitor identity, or *sox19a* but not *pcna*, indicative of neural precursors (Fig. 2G,H). Notably, whereas progenitors and precursors closely aligned with differentiating neurons expressed *neurog1*, which encodes a pro-neuronal transcription factor (Cau et al., 1997; Kim et al., 1997), progenitors more closely related to oligodendrocyte lineage cells (marked by expression of *olig2* and *sox10*) did not express *neurog1* (Fig. 2E-J). These data suggest that pMN cells include two different progenitor populations: *sox19a*^+^ *neurog1*^+^ pMN neuronal progenitors and *sox19a*^+^ *neurog1*^−^ Pre-OPC progenitors.

### OPCs and pre-myelinating oligodendrocytes form distinct clusters at 48 hpf

To begin teasing apart the origins of pMN progenitor population heterogeneity in the zebrafish spinal cord, we used unsupervised Seurat clustering to identify temporally specific cell populations within each collected time point. We again determined cluster identities and unique markers by top differentially expressed genes within each population (Figs. 2-5; Figs. S3-S5).

At 48 hpf the majority of the cells were *olig2*^*–*^ post-mitotic neurons with few *olig2*^*+*^ progenitor cells and glia (Fig. 3A-C). Most of the *elavl4*^*+*^ neurons (Fig. 3D) consisted of *isl1*^*+*^ *chodl*^*+*^ *mnx1*^*+*^ motor neurons (Enjin et al., 2010; Ericson et al., 1992; William et al., 2003) (Fig. S3D,E,G). There were two progenitor clusters we identified as *sox19a*^*+*^ *pcna*^*+*^ *gfap*^*–*^ pMN1 cells and *sox19a*^*+*^ *pcna*^*–*^ *gfap*^*+*^ radial glia (RG) (Fig. 3E,F; Fig. S3C). RG persist as dividing cells into larval and adult stages and can produce new neurons following spinal cord injury (Briona and Dorsky, 2014; Johnson et al., 2014; Johnson et al., 2016; Park et al., 2007). Three clusters of 48 hpf cells expressed *sox10*, a canonical marker of oligodendrocyte lineage cells (Fig. 3G) (Britsch et al., 2001; Kuhlbrodt et al., 1998; Park et al., 2002). One group of cells expressed *foxd3* (Fig. S3F) identifying these cells as MEP glia, which emerge from the ventral spinal cord at motor exit points (Fontenas and Kucenas, 2021; Kucenas et al., 2008a; Smith et al., 2014). A second group of *sox10*^*+*^ cells expressed *sox5, sox9a, sox9b* and *olig1*, them as OPCs (Baroti et al., 2016; Schebesta and Serluca, 2009; Stolt et al., 2006; Zhou and Anderson, 2002) (Fig. 3H-K), whereas the third cluster expressed *tcf7l2* and *myrf* (Fig. 3L,M), indicating a population of pre-myelinating oligodendrocytes (Pre-OLs) (Emery et al., 2009; Hammond et al., 2015). Consistent with the integrated dataset, oligodendrocyte lineage cells were separated from the neuronal lineage denoted by differential *neurog1* expression (Fig. 3N). Further, cells with gene expression characteristic of Pre-OPCs (*sox19a*^+^ *neurog1*^−^) were absent, indicating that these cells must arise earlier in development but are then depleted following OPC formation.

**Fig 3.**
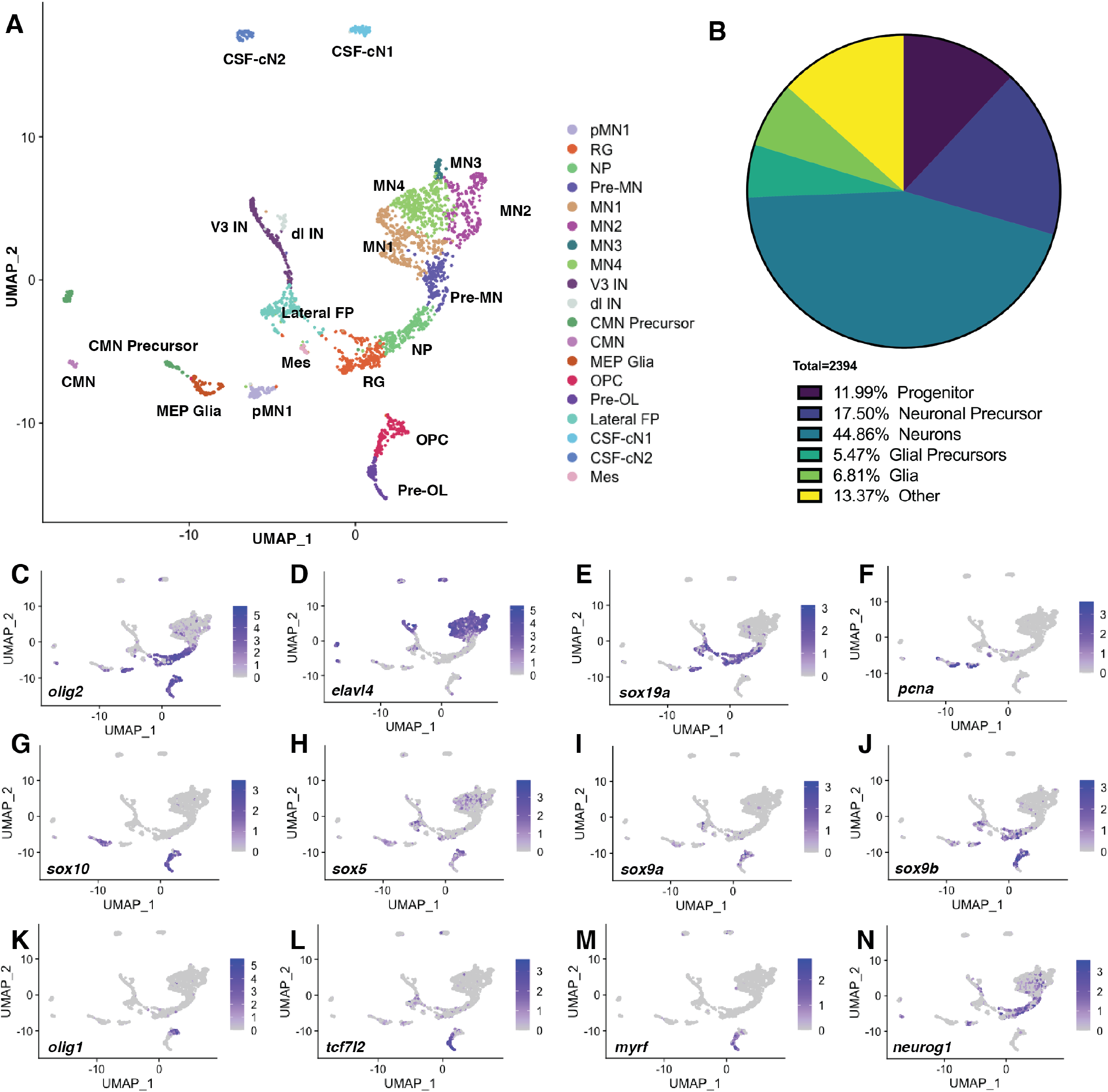
Cell type clustering and markers of pMN cells at 48 hpf. (A) Unsupervised UMAP of scRNA-seq data from *olig2:*EGFP^+^ spinal cord cells obtained from 48 hpf *Tg(olig2:EGFP)* embryos identifies 19 clusters. Each point represents one cell (n=2394). (B) Proportions of pMN gross cell type identities at 48 hpf. A majority of isolated pMN cells are post-mitotic neurons; ∼12% are progenitors and ∼12% are glial lineages. (C-N) UMAP feature plots of selected transcripts. Cells are colored by expression level (gray is low, purple is high). o*lig2*^*–*^ *elav4*^*+*^ cells represent post-mitotic neuronal populations (C,D). Co-expression of *olig2, sox19a* and *pcna* designate a small population of pMN progenitors and radial glia (RG) (E,F). *sox10* expression signifies MEP glia and the oligodendrocyte lineage (G). Expression of *sox5, sox9a/b* and *olig1* represent OPCs (H-K), whereas *tcf7l2* and *myrf* expression mark Pre-OLs. (L-M). *neurog1* represents neuronal progenitors and precursors (N).

### pMN cells express genes characteristic of Pre-OPCs at 24 and 36 hpf

To identify progenitors that initially give rise to the oligodendrocyte lineage we assessed populations of pMN cells at 36 and 24 hpf (Figs. 4 and 5). Our 36 hpf dataset included fewer *elavl4*^*+*^ differentiated neurons and more presumptive progenitors and neuronal precursors than the 48 hpf dataset (Fig. 4A-C). In addition to pMN1 progenitors, we identified distinct *olig2*^+^ *sox19a*^+^ *neurog1*^+^ progenitor clusters that we labeled as pMN2 and pMN3 (Fig. 4A,D-F). pMN2 cells were marked by relatively high expression of *plk1* and *aspm*, indicative of cells undergoing mitosis (M-Phase), whereas pMN3 cells had relatively high levels of *hells* expression characteristic of DNA replication in S-Phase (Avides and Glover, 1999; Barr et al., 2004; Fish et al., 2006; Mjelle et al., 2015) (Figs. S4 and S5C,D). RGs were not evident, indicating that cells with radial glia characteristics emerge between 36 and 48 hpf. Notably, few cells expressed appreciable levels of *sox10* at 36 hpf, suggesting that OPCs were not yet specified (Fig. S4). However, we identified a cluster consisting of *sox19a*^*+*^ *olig2*^*+*^ cells that also expressed *nkx2*.*2a*, most of which were *neurog1*^*–*^ (Fig. 4D-G). Spinal cord OPCs arise from *olig2*^*+*^ *nkx2*.*2a*^*+*^ progenitors (Agius et al., 2004; Fu et al., 2002; Kessaris et al., 2001; Kucenas et al., 2008b; Soula et al., 2001; Zhou et al., 2001) raising the possibility that this population is the source of OPCs. Consistent with this possibility, a subset of these cells also expressed low levels of *olig1* (Fig. 4H). These cells also expressed genes that have been used to characterize a Pre-OPC cell type, including *ascl1a* and *gsx2* (Fig. 4I,J) (Parras et al., 2007; Sugimori et al., 2007; Zhang et al., 2020). Therefore, *ascl1a*^*+*^ *gsx2*^*+*^ cells may represent a population of pMN cells transitioning to OPCs at 36 hpf.

**Fig 4.**
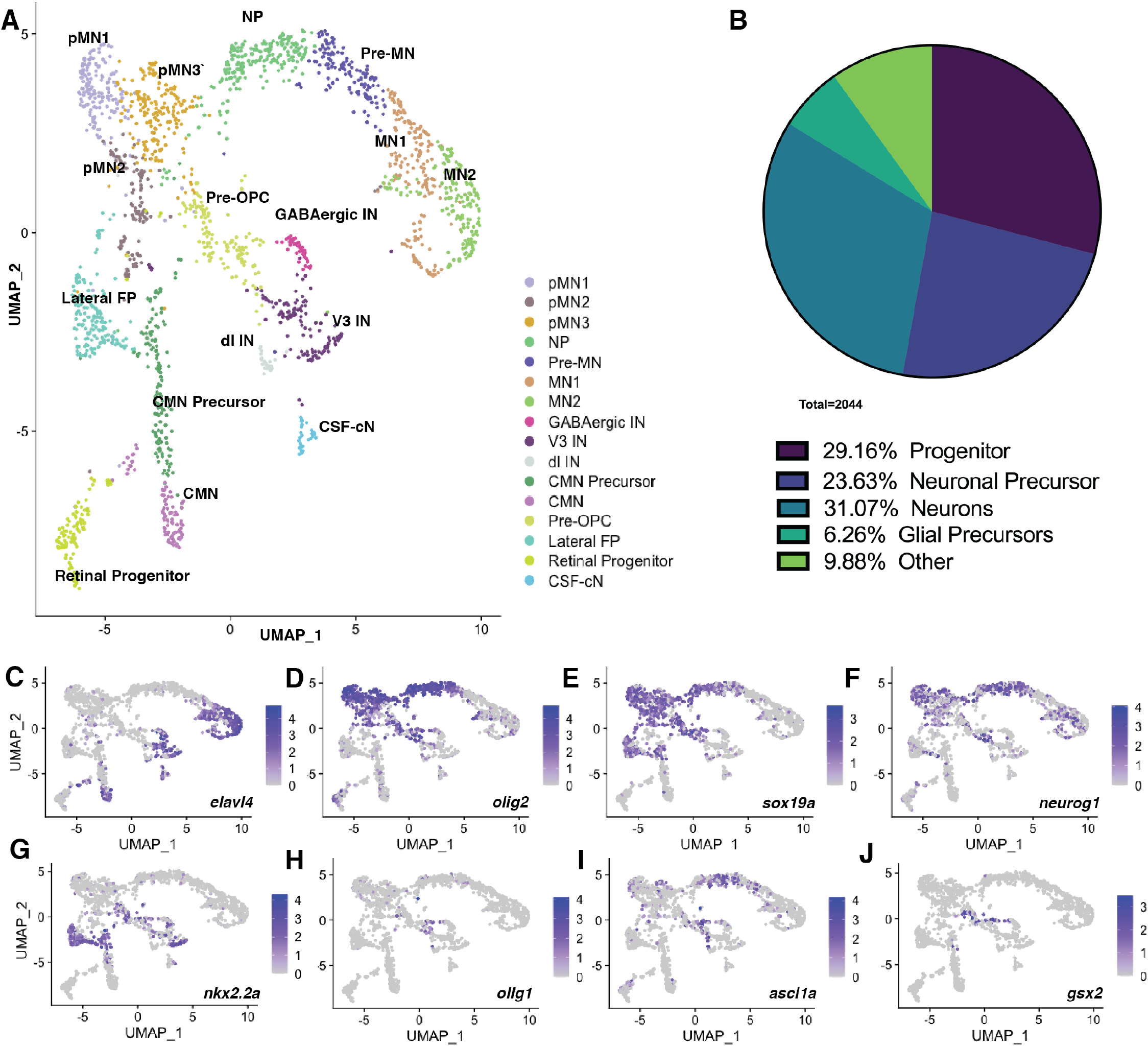
Cell type clustering and markers of pMN cells at 36 hpf. (A) Unsupervised UMAP of scRNA-seq data from *olig2:*EGFP^+^ spinal cord cells obtained from 36 hpf *Tg(olig2:EGFP)* embryos identifies 16 clusters. Each point represents one cell (n=2044). (B) Proportions of pMN gross cell type identities at 36 hpf. A majority of isolated pMN cells are neurons or neuronal precursors; ∼30% are progenitors and ∼6% are glial precursors. (C-F) UMAP feature plots of selected transcripts. Cells are colored by expression level (gray is low, purple is high). (C-J) UMAP feature plots of selected transcripts. Cells are colored by expression level (gray is low, purple is high). *elav4*^*+*^ cells represent post-mitotic neuronal populations (C). Co-expression of *olig2* and *sox19a* designate pMN progenitors (D,E) and expression of *neurog1* in a subset of *olig2*^*+*^ *sox19*^*+*^ cells represent neuronal progenitors and precursors (F). Combinatorial expression of *nkx2*.*2a, olig1, ascl1a, gsx2* and *her4*.*2* likely represent an OPC progenitor population (G-J).

**Fig 5.**
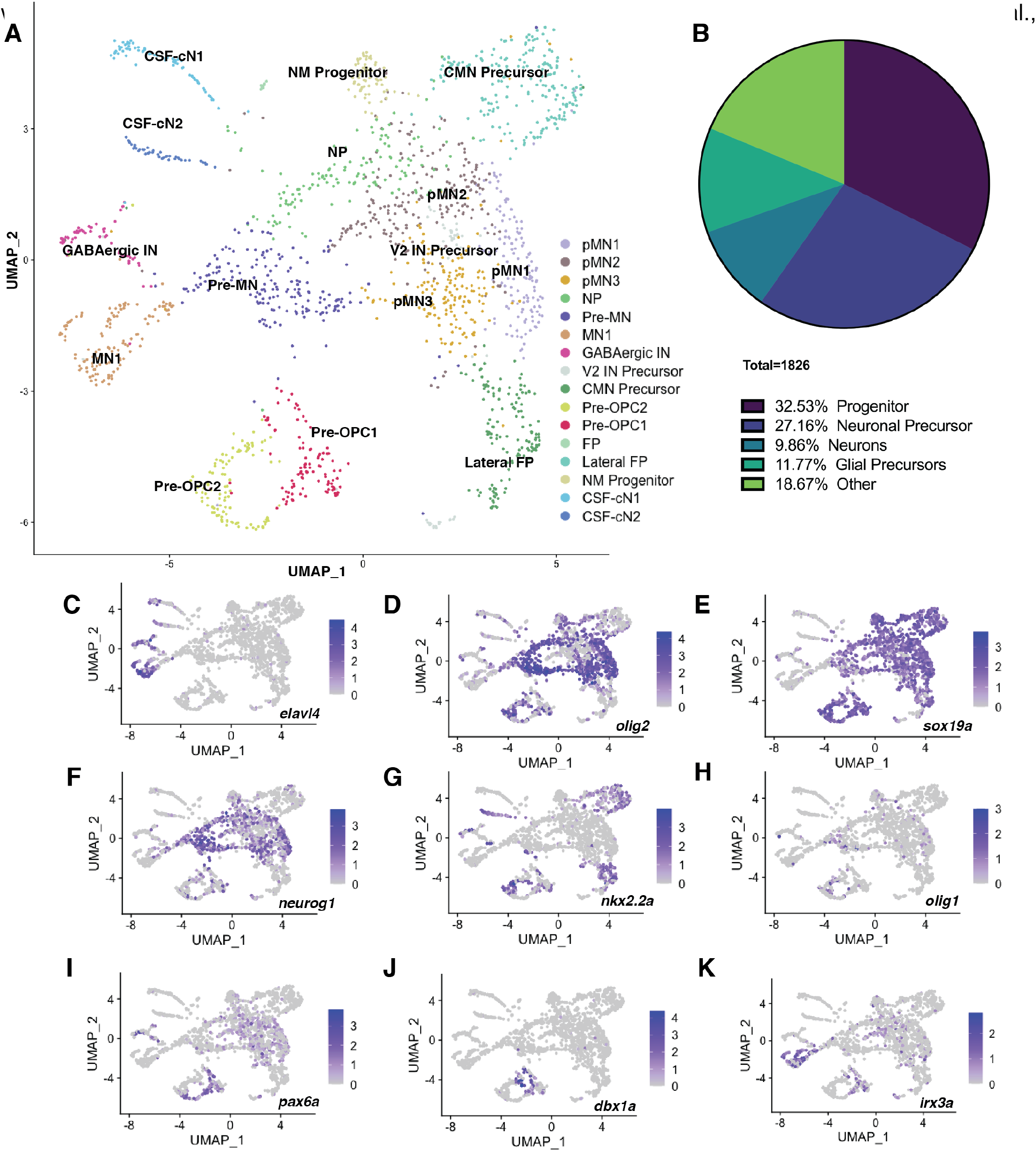
Cell type clustering and markers of pMN cells at 24 hpf. (A) Unsupervised UMAP of scRNA-seq data from *olig2:*EGFP^+^ spinal cord cells obtained from 24 hpf *Tg(olig2:EGFP)* embryos identifies 16 clusters. Each point represents one cell (n=1826). (B) Proportions of pMN gross cell type identities at 24 hpf. A majority of isolated pMN cells are progenitors or precursors whereas ∼10% are neurons and ∼12% are glial precursors. (C-K) UMAP feature plots of selected transcripts. Cells are colored by expression level (gray is low, purple is high). *elav4*^*+*^ cells represent post-mitotic neuronal populations (C). Co-expression of *olig2* and *sox19a* designate pMN progenitors (D,E) and expression of *neurog1* in subsets of *olig2*^*+*^ *sox19*^*+*^ cells represent neuronal progenitors and precursors (F). Expression of *olig2, nkx2*.*2a* and *olig1* represent the more mature Pre-OPC2 population (D,G,H) and lower *olig2* expressing cells that are also marked by *pax6a, dbx1a* and *irx3a* represent a population of less mature Pre-OPC1 cells (I-K).

Cell clusters within the 24 hpf dataset were analogous to the 36 hpf dataset, consisting of neuronal progenitors, committed neuronal precursors and differentiating motor neurons and interneurons (Fig. 5A-E). Similar to the 36 hpf data, we identified cells with characteristics of Pre-OPCs, but these were subdivided into two clusters that we designated as Pre-OPC1 and Pre-OPC2 (Fig. 5A,D-K). Pre-OPC1 cells expressed low levels of *olig2, olig1* and *nkx2*.*2* but higher levels of genes associated 1992) (Fig. 5G-K). By contrast, the Pre-OPC2 population expressed higher levels of *olig2, olig1* and *nkx2*.*2* but lower levels of *pax6a, dbx1a* and *irx3a* (Fig. 5G-K). One possible explanation for this clustering is that Pre-OPCs progress from a less mature Pre-OPC1 state to a more mature Pre-OPC2 state. To test this possibility, we used RNA velocity analysis (Bergen et al., 2020; La Manno et al., 2018) to predict the trajectories of cells within the 24 hpf dataset. These data show that a majority of Pre-OPC1 cellular velocities are directed towards the Pre-OPC2 population (Fig. S6), supporting the prediction that Pre-OPC1 cells transition to Pre-OPC2 cells. Altogether, these data suggest that OPCs arise from a dedicated Pre-OPC progenitor population that exists during active neurogenesis at 24 hpf.

### Motor neurons and oligodendrocytes have separate developmental trajectories throughout pMN development

Our data raise the possibility that pMN cells include distinct subpopulations of pMN progenitors, one population that produces neurons and a separate population of Pre-OPCs that later form oligodendrocytes. To further investigate the relationships between these specific progenitor populations and their derivative cell types, we employed RNA velocity analysis to predict lineage trajectories within the integrated dataset (Fig. S7). The projection of velocity fields onto the integrated UMAP and single-cell plotting of RNA velocity vector fields highlighted separate developmental trajectories for several pMN cell populations including glial, motor neuron, CMN, and interneuron lineages (Fig. S7). The major predicted trajectory consisted of pMN1-3 progenitors transitioning into neural precursors (NP), which differentiate into motor neurons.

Separate from the neuronal trajectory, RNA velocity analysis predicted that the Pre-OPC population consists of two detached populations with one vector field transitioning into OPCs and the other into the V3 interneuron population (Fig. S7A). To better delineate the trajectories observed branching from the Pre-OPC cluster, we assessed the RNA velocity vector field at a single-cell level, allowing us to evaluate projected commitment of each cell based on individual vector size and directionality. This revealed that approximately half of the cells in the Pre-OPC lineage had larger vectors with strong directionality towards the interneuron cluster, suggestive of an interneuron precursor population. The other portion of cells within the Pre-OPC population had smaller vectors with less directionality, denoting a less committed progenitor identity. A majority of these trajectories pointed toward the oligodendrocyte lineage and those closest to OPCs had larger vectors with stronger directionality, indicative of cells undergoing specification for OPC fate (Fig. S7B). These data indicate that the integrated Pre-OPC cluster may be composed of interneuron precursors and Pre-OPCs.

To further resolve these distinct cell types within the Pre-OPC cluster, we subsetted and sub-clustered the oligodendrocyte lineage. The Pre-OPC population was split into three sub-clusters encompassing cells collected from 24 and 36 hpf samples (Fig. S8A-B). Pre-OPC1/2 mainly consisted of cells in G2M/S phase, suggestive of a more progenitor-like identity, consistent with the small and less directional velocities of these cells. By contrast, Pre-OPC3 cells were in G1, thereby aligning with the preceding velocity analysis where these cells had larger and more directional vectors, indicative of committed cells (Fig. S8C). Consistent with this idea, we found that cells within the Pre-OPC3 sub-cluster expressed transcripts characteristic of neuronal precursors, including *neurod4, neurog1* and *dlx2a* (Cau et al., 1997; Farah et al., 2000; Kim et al., 1997; Petryniak et al., 2007) whereas Pre-OPC1/2 cells did not (Fig. S8D-F). Moreover, cells within Pre-OPC1/2 expressed *nkx2*.*2a* at a higher level than cells in the Pre-OPC3 cluster, in line with our prediction that these cells are fated to the oligodendrocyte lineage (Fig. S8G). One possible explanation for why these two populations were clustered together could be based on their shared expression of unique transcripts like *barhl2* and *otx2a* (Fig. S8H-K). This split in the Pre-OPC lineage led us to remove the Pre-OPC3 sub-cluster from the oligodendrocyte lineage before continuing further pseudotime analysis (Fig. S8L).

Our evaluation of RNA velocity fields implied that motor neurons and oligodendrocytes arise from discrete progenitor populations. However, the exact progression of these lineages and cellular position through time is not discernable in this analysis. To gain a deeper understanding of the differences in distinct lineages and temporal ordering within the motor neuron and oligodendrocyte lineages, we used Slingshot (Street et al., 2018) as an additional trajectory analysis. We performed Slingshot analysis on subsetted clusters included in the oligodendrocyte or motor neuron lineages (Fig. 6A; Fig. S9A). Slingshot predicted four lineages within the neuronal subset, all emerging from pMN1. The first three lineages progressed through pMN2/3, NP1/2, Pre-MN and MN1. However, the first lineage continued through MN4 ending at MN5, the second went through MN4 stopping at MN2 and the third lineage terminated at MN3 (Fig. S9A). The fourth predicted lineage within the neuronal trajectory progressed through pMN2/3 and terminated at RG (Fig. S9A). These data are consistent with differences in motor neuron subtype formation and the creation of a radial glial population during spinal cord development (Mizuguchi et al., 2001; Park et al., 2007; Stifani, 2014). For the subsetted oligodendrocyte lineage clusters, Slingshot predicted only one lineage that began at Pre-OPC1, transitioned through Pre-OPC2, OPC1/2 and ended at Pre-OLs (Fig. 6A). When we aligned the oligodendrocyte and second predicted neuronal Slingshot lineages along pseudotime, we found that Pre-OPC1 and pMN1 progenitors represented the least differentiated cell types within the oligodendrocytes and neuronal lineages, respectively (Fig. 6B; Fig. S9B). These results reinforce our conclusion that oligodendrocytes and motor neurons arise from two distinct populations of progenitors.

**Fig 6.**
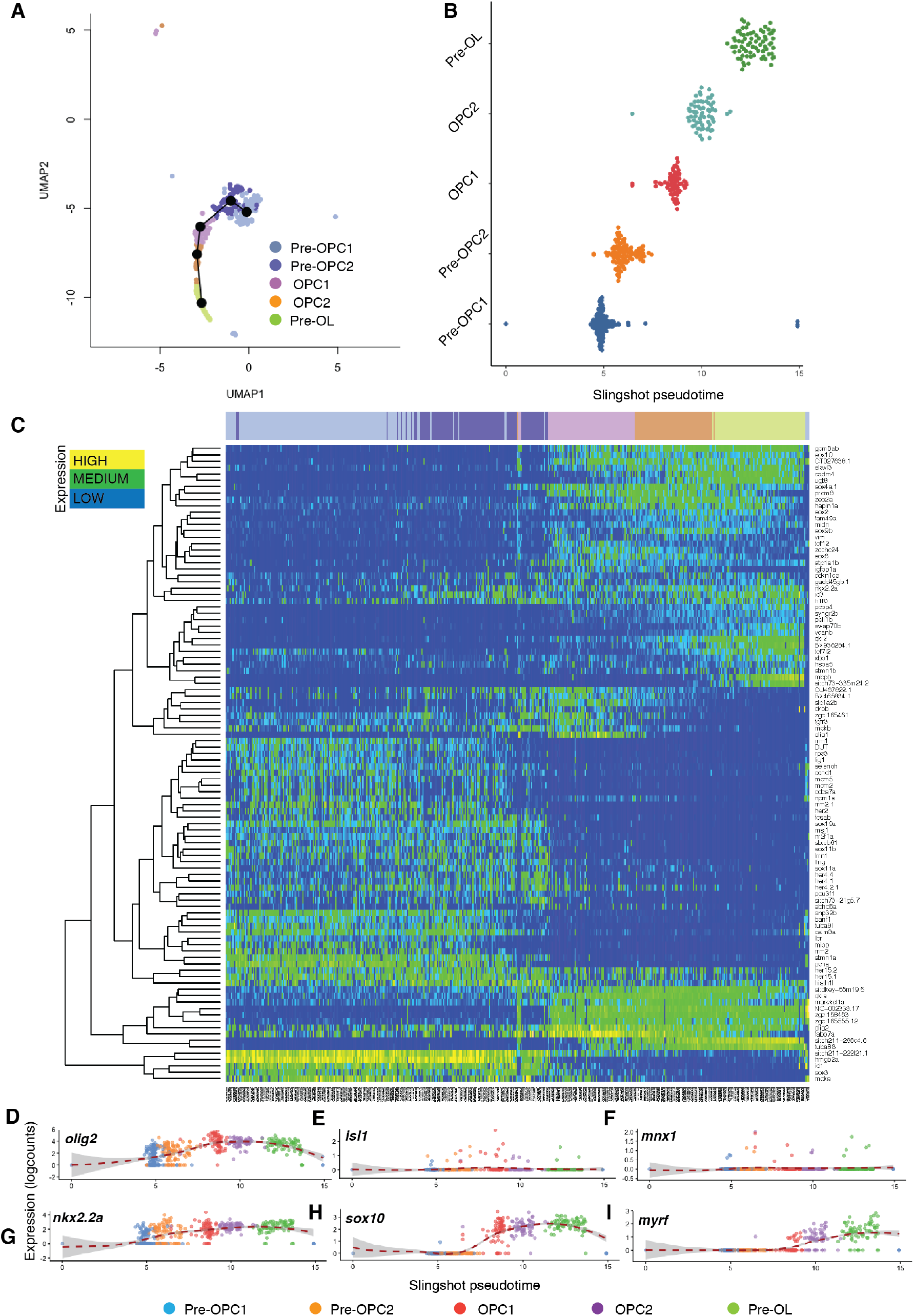
Pseudotime analysis of pMN cells reveals gene expression characteristic of oligodendrocyte development. (A) UMAP of sub-clustered oligodendrocyte lineage cells from the integrated dataset depicting the Slingshot-predicted lineage trajectory. (B) Depiction of cells within each sub-cluster from the Slingshot-predicted lineage along pseudo-temporal ordering. Each dot represents a cell and its predicted temporal position in development. (C) Heatmap showing top 100 differentially expressed transcripts based on the Slingshot-predicted oligodendrocyte lineage (left = OPC Progenitors, right = Pre-OLs). Dendrogram displays hierarchical clustering; each column represents the relative expression of each labeled transcript in a single-cell. (D-I) Scater smooth expression plots for transcripts related to neuronal (D-F) or oligodendrocyte (F-I) development within each sub-cluster along Slingshot pseudo-temporal ordering. Each point represents the expression of each transcript within a single-cell. Most of the oligodendrocyte lineage cells do not express neuronal transcripts *isl1* and *mnx1* (E,F). *olig2* and *nkx2*.*2a* are expressed in Pre-OPCs and their expression is maintained throughout the lineage (D,G). OPCs and Pre-OLs are marked by *sox10* expression (H). OPC2 and Pre-OLs are designated by increased expression of *myrf* (I).

The separated neuronal and glial pMN trajectories implies that there are differences in gene expression that guide motor neuron and oligodendrocyte developmental progression. To explore these potential differences, we fit a generalized additive model (GAM) to each predicted lineage to evaluate the top 100 differentially expressed genes related with pseudotime (Fig. 6C; Fig. S9C). In the neuronal lineage, genes such as *isl1, mnx1, dla* and *neurod4* were present among the top differentially expressed genes, accordant with known transcripts required for motor neuron formation (Fig. S9C) (Appel and Eisen, 1998; Appel et al., 2001; Arber et al., 1999; Ericson et al., 1992; Farah et al., 2000; Tanabe et al., 1998). Comparatively, genes associated with oligodendrocyte development like *sox9b, prdm8, olig1* and *nkx2*.*2a* were within the differentially expressed genes in the oligodendrocyte lineage but not the neuronal lineage (Fig. 6C; Fig. S9C) (Agius et al., 2004; Kessaris et al., 2001; Schebesta and Serluca, 2009; Scott et al., 2020; Soula et al., 2001; Stolt et al., 2003; Stolt et al., 2006; Zhou and Anderson, 2002; Zhou et al., 2001).

We then used smoother plots overlaid onto single-cell expression through pseudotime of select transcripts associated with motor neuron or oligodendrocyte development to evaluate the biological accuracy of our Slingshot analysis (Fig. 6D-I; Fig. S9D-I). In agreement with known developmental progression, progenitors and precursors in the neuronal lineage had high expression of *olig2* but cells in MN1-4 expressed it at low levels (Fig. S9D-I) (Novitch et al., 2001). By contrast, oligodendrocyte lineage cells maintained *olig2* expression at high levels through all stages (Fig. 6D) (Zhou et al., 2000; Zhou et al., 2001). Consistent with motor neuron development, *isl1* expression began in NP1/2 and *mnx1* and *isl1* expression were upregulated and maintained in motor neuron populations (Fig. S9E-F) (Arber et al., 1999; Ericson et al., 1992). Conversely, very few cells in the oligodendrocyte lineage expressed *isl1* or *mnx1* (Fig. 6E-F). *nkx2*.*2a* was expressed throughout the oligodendrocyte lineage, *sox10* expression was evident in OPC1/2 and maintained in Pre-OLs, and most cells within the OPC2 and Pre-OL cluster expressed *myrf* (Fig. 6G-I). A small number of cells in pMN1/2 expressed *nkx2*.*2a* and *sox10* and almost no cells within the neuronal lineage expressed *myrf* (Fig. S9G-I). Overall, the lineage trajectory predictions support the hypothesis that oligodendrocytes and motor neurons arise from distinct progenitors and the subsequent gene expression analyses offer evidence for the accuracy of the Slingshot predicted lineages.

### Pre-OPCs are formed prior to OPC specification in the embryonic zebrafish spinal cord

Thus far we have provided evidence for the existence of discrete progenitors within the pMN lineage within our scRNA-seq datasets. To assess the temporal development of these two populations, we compared the expression of *ascl1a, neurog1* and *gsx2* within the predicted lineages along pseudotime with smoother plots overlaid onto single-cell expression of each gene (Fig. 7A-C). Expression of *ascl1a* was highest in the NP1/2 populations and in a subset of Pre-OPCs but was downregulated in differentiated populations within both lineages (Fig. 7A). Most strikingly, *gsx2* was expressed exclusively in Pre-OPC1/2 and *neurog1* was mainly expressed within the progenitors and precursors of the neuronal lineage (Fig. 7B-C). Based on these data we expect that Pre-OPCs (*gsx2*^*+*^*/neurog1*^*–*^) can be distinguished from the pMN population (*gsx2*^*–*^*/neurog1*^*+*^) in vivo by assessing differential expression of *gsx2* and *neurog1*.

**Fig 7.**
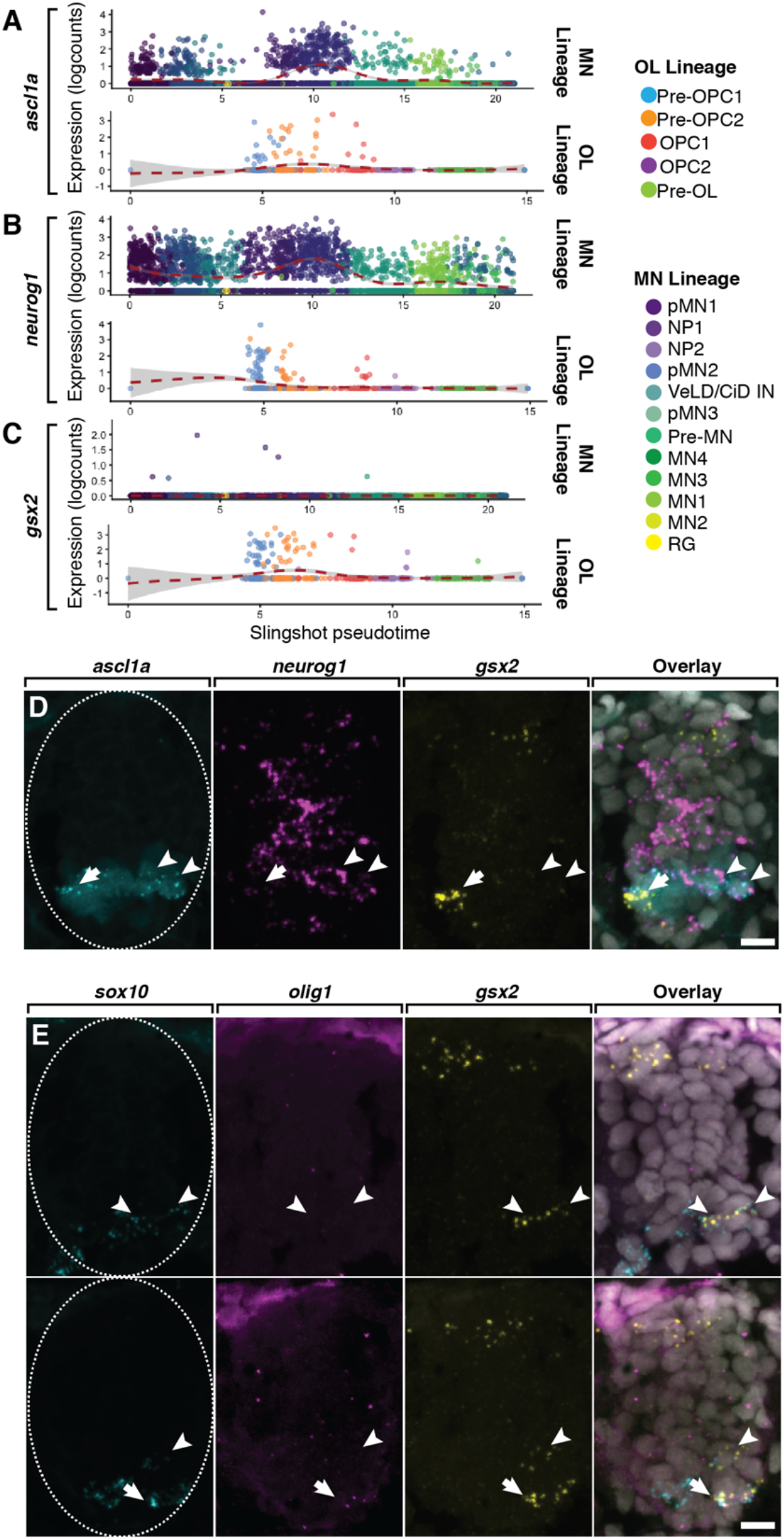
In vivo identification of distinct neuronal and oligodendrocyte progenitors in the embryonic zebrafish spinal cord. (A-C) Scater smooth expression plots for transcripts distinguishing pMN progenitors and Pre-OPCs within each lineage along Slingshot pseudo-temporal ordering. Each point represents the expression of each transcript within a single-cell. A portion of cells in the neuronal and oligodendrocyte lineages express *ascl1a* (A). *neurog1* is highly expressed by many cells in the neuronal lineage but expressed by relatively few cells in the oligodendrocyte lineage (B). *gsx2* is exclusively expressed in the Pre-OPC population (C). (D,E) Transverse trunk spinal cord sections obtained from zebrafish embryos processed for fluorescent RNA ISH. Dorsal is up. (D) Detection of *ascl1a* (cyan puncta), *neurog1* (magenta puncta), and *gsx2* (yellow puncta) mRNA in 36 hpf *Tg(olig2:EGFP)* embryos to label pMN cells (light cyan cell bodies). Arrowheads denote *ascl1a*^*+*^*/neurog1*^*+*^*/gsx2*^*–*^ neuronal pMN progenitors. Arrow highlights an *ascl1a*^*+*^*/neurog1*^*–*^*/gsx2*^*+*^ Pre-OPC. (E) Detection of *sox10* (cyan puncta), *neurog1* (magenta puncta), and *gsx2* (yellow puncta) mRNA in 40 hpf embryos. Arrowheads denote *sox10*^*+*^*/olig1*^*–*^*/gsx2*^*+*^ Pre-OPCs and the arrow highlights a *sox10*^*+*^*/olig1*^*+*^*/gsx2*^*+*^ Pre-OPC. Dashed oval outlines the spinal cord boundary.

To validate the separation of Pre-OPCs and neuronal-fated pMN progenitor populations, we performed fluorescent in situ hybridization (FISH) probing for *ascl1a, gsx2* and *neurog1* mRNA at 36 hpf in *Tg(olig2:EGFP)* embryos. Consistent with our scRNA-seq data, both *ascl1a*^*+*^ *neurog1*^*+*^ *gsx2*^*–*^ cells and *ascl1a*^*+*^ *neurog1*^*+*^ *gsx2*^*+*^ cells were evident in the ventral spinal cord (Fig. 7D). We next tested the prediction that *gsx2* expression marks cells fated to develop as OPCs by probing for *sox10, olig1* and *gsx2* transcripts at 40 hpf. This revealed that a majority of *sox10*^*+*^ cells expressed *gsx2* and a subset of those cells also expressed *olig1* (Fig. 7E). These data are consistent with predictions made from our scRNA-seq data, thereby revealing transcriptionally distinct spinal cord progenitors of ventral neurons and oligodendrocytes.

## Discussion

In the developing nervous system, neural progenitor populations first give rise to neurons and then glia, a process guided by changes in environmental cues and alterations in gene expression (Kessaris et al., 2001; Miller and Gauthier, 2007). Though the generation of diverse neurons and glia are imperative to nervous system function, the origins of neuronal and glial precursors remain elusive. Here we provide evidence that in the ventral spinal cord, motor neurons and oligodendrocytes arise from distinct neural progenitor populations that are specified early in development.

The major finding revealed by our scRNA-seq dataset and corroborated by in vivo mRNA expression analysis was the presence of two transcriptionally distinct progenitor populations. One population of cells highly expressed the proneuronal transcript *neurog1* and was closely associated with the motor neuron lineage. By contrast, a majority of cells in the other progenitor population did not express *neurog1* and were closely related to the oligodendrocyte lineage, suggesting that motor neurons and oligodendrocytes arise from independent progenitors. Consistent with our findings, most clonally related cells in the mouse cortex consist of either neurons or glia (Luskin et al., 1988; Luskin et al., 1993; McCarthy et al., 2001). Similarly, through time-lapse imaging of transgenic labeled pMN cells our group showed that motor neurons and OPCs arise from distinct pMN progenitor lineages (Ravanelli and Appel, 2015). Further, pMN progenitors isolated during oligodendrogenesis and transplanted into chick embryos during motor neuron formation were only able to form oligodendrocytes (Mukouyama et al., 2006), suggesting that a subset of pMN progenitors are restricted for oligodendrocyte fate.

Our pseudotime lineage analysis provided further support for the distinction between motor neuron and oligodendrocyte lineages by affirming that transcripts expressed throughout the predicted trajectories for motor neurons and oligodendrocytes reflected established developmental mechanisms. For example, cells identified as having motor neuron fates expressed *isl1*, which encodes a LIM homeodomain transcription factor required for motor neuron survival and specification (Ericson et al., 1992; Pfaff et al., 1996), whereas cells in the oligodendrocyte lineage did not express *isl1*. In comparison, OPCs and differentiating oligodendrocytes highly expressed the transcript *sox10*, a gene that encodes a transcription factor necessary for oligodendrocyte formation and maintenance (Britsch et al., 2001; Kuhlbrodt et al., 1998; Park et al., 2002; Takada et al., 2010). Further, trajectory analysis showed that *prdm8* is expressed in OPCs and downregulated in Pre-OLs, consistent with our recent publication in which we found that *prdm8* inhibits oligodendrocyte differentiation (Scott et al., 2020). These data indicate that motor neurons and oligodendrocyte arise from distinct progenitors that undergo independent developmental progression and transcriptional alterations.

Through further evaluation of the differences between pMN progenitors and Pre-OPCs we identified a novel expression profile that can be used to explore the development of the oligodendrocyte lineage in future studies. The majority of Pre-OPCs did not express *neurog1*, but instead uniquely expressed *gsx2*. The mutually exclusive expression of *gsx2* and *neurog1* in pMN progenitors is similar to previously characterized expression of these transcripts in zebrafish dorsal spinal cord progenitors (Satou et al., 2013). As Pre-OPCs undergo specification and subsequent differentiation into oligodendrocytes, *gsx2* expression is down regulated. The downregulation of *gsx2* by oligodendrocyte lineage cells is consistent with previous findings that loss of Gsx2 function in the mouse cortex resulted in precocious OPC formation at the expense of neurons, indicating that Gsx2 restricts OPC formation and differentiation (Chapman et al., 2012; Chapman et al., 2018). Therefore, it is possible that Gsx2 functions to reserve a population of pMN progenitors as Pre-OPCs during motor neuron formation and is later downregulated at the onset of gliogenesis to allow for OPC specification.

We also found that at 24 hpf Pre-OPCs were composed of two clusters, a less mature Pre-OPC1 population and a more mature Pre-OPC2 population. Of note, Pre-OPC1 cells expressed *olig2* at a lower level compared to Pre-OPC2 cells and additionally expressed transcripts characteristic of more dorsal progenitors such as *dbx1a* and *pax6a* (Briscoe et al., 2000; Lu et al., 1992). These expression data are compatible with previous findings by our lab that motor neurons and OPCs arise from distinct progenitors that successively initiate expression of *olig2* as more dorsal progenitors slide ventrally into the pMN domain (Ravanelli and Appel, 2015; Ravanelli et al., 2018). Therefore, the Pre-OPC1 cluster we identified from our scRNA-seq analysis might represent a transient population of dorsally recruited progenitors that initiate *olig2* expression while retaining transcripts characteristic of their more dorsal spinal cord origin.

## METHODS AND MATERIALS

### Zebrafish lines and husbandry

All animal work was approved by the Institutional Animal Care and Use Committee (IAUCUC) at the University of Colorado School of Medicine. All non-transgenic embryos were obtained from pairwise crosses of males and females from the AB strain. Embryos were raised at 28.5°C in E3 media (5 mM NaCl, 0.17 mM KCl, 0.33 mM CaCl2, 0.33 mM MgSO4 at pH 7.4, with sodium bicarbonate), sorted for good health and staged accordingly to developmental morphological features and hours post-fertilization (hpf) (Kimmel et al., 1995). Developmental stages are described in the results section for individual experiments. Sex cannot be determined at embryonic and larval stages. The transgenic line used was *Tg(olig2:EGFP)*^*vu12*^ (Shin et al., 2003). All transgenic embryos were obtained from pairwise crosses of males or females from the AB strain to males or females to the *Tg(olig2:EGFP)*^*vu12*^ strain.

### Fluorescent In situ RNA Hybridization

Fluorescent in situ RNA hybridization was performed using the RNAScope Multiplex Fluorescent V2 Assay Kit (Advanced Cell Diagnostics; ACD) on 12 μm thick paraformaldehyde-fixed and agarose embedded cryosections according to manufacturer’s instructions with the following modification: embryos were not dehydrated prior to embedding and slides were covered with parafilm for all 40°C incubations to maintain moisture and disperse reagents across the sections. The zebrafish *ascl1a-C1, neurog1*-C2, and *gsx2*-C3 transcript probes were designed and synthesized by the manufacturer and used at 1:50 dilutions. Transcripts were fluorescently labeled with Opal520 (1:1500), Opal570 (1:500) and Opal650 (1:1500) using the Opal 7 Kit (NEL797001KT; PerkinElmer).

### Imaging

Fixed sections of embryos were imaged on a CellObserver SD 25 spinning disk confocal system (Carl Zeiss). RNA FISH images were acquired using a 40X (n.a. 0.75) objective. Images are reported as extended z-projections collected using Zen (Carl Zeiss) imaging software. Image brightness and contrast were adjusted in Photoshop (Adobe) or ImageJ (National Institutes of Health).

### Cell dissociation and FACs

Methods for cell dissociation and sorting were previously reported (Scott et al.). 24, 36, and 48 hpf *Tg(olig2:EGFP)* euthanized embryos were collected in 1.7 ml microcentrifuge tubes and deyolked in 100 μl of pre-chilled Ca free Ringers solution (116 mM NaCl, 2.6 mM KCl, 5 mM HEPES, pH 7.0) on ice. Embryos were pipetted intermittently with a P200 micropipettor for 15 minutes and incubated unperturbed for 5 min. 500 μl of protease solution (10 mg/ml BI protease, 125 U/ml DNase, 2.5 mM EDTA, 1X PBS) was added to microcentrifuge tubes on ice for 15 min and embryos were homogenized every 3 min with a P100 micropipettor for 15 min. 200 μl of STOP solution (30% FBS, 0.8 mM CaCl2, 1X PBS) was then mixed into the tubes. Samples were then spun down at 400g for 5 min at 4°C and supernatant was removed. On ice, 1 ml of chilled suspension media (1% FBS, 0.8 mM CaCl_2_, 50 U/ml Penicillin, 0.05 mg/ml Streptomycin) was added to samples and then spun down again at 400g for 5 min at 4°C. Supernatant was removed, and 400 μl of chilled suspension media was added and solution was filtered through a 35 μm strainer into a collection tube. Cells were FAC sorted to distinguish EGFP^+^ cells using a MoFlo XDP100 cell sorter at the CU-SOM Cancer Center Flow Cytometry Shared Resource and collected in 1.7 ml FBS coated microcentrifuge tubes in 200 μl of 1X PBS.

### Single-cell cDNA library preparation and raw data generation

The library preparation and sequencing to produce the data reporter here were described previously (Scott et al., 2020). The Chromium Box from 10X Genomics was used to capture cells using Chromium Single Cell 3’ Reagent Kit part no. PN-1000075. Libraries were sequenced on the Illumina NovaSEQ6000 Instrument. FASTQ files were analyzed using Cell Ranger Software. 2174 (24 hpf), 2555 (36 hpf) and 3177 (48 hpf) cells were obtained yielding a mean of 118,014 (24 hpf), 65,182 (36 hpf) and 96,053 (48 hpf) reads per cell with a median of 1929 (24 hpf), 1229 (36 hpf) and 1699 (48 hpf) genes identified per cell. Raw sequencing reads were demultiplexed, mapped to the zebrafish reference genome (build GRCz11/danRer11) and summarized into gene expression matrices using CellRanger (version 3.0.1).

### Unsupervised Seurat analysis of individual stages

Resulting count matrices for each development time point were loaded into Seurat (v3.2.1) and mitochondrial gene expression was added by PercentageFeatureSet (pattern = “^mt-”). Cell barcodes with fewer than 200 detectable genes, more than 5% of nFeature_RNAs derived from mitochondrial genes, or more than 50,000 nFeature_RNAs (24 hpf, 48 hpf) and 30,000 UMIs (36 hpf) were removed, resulting in 1952, 2149 and 2393 cells filtered for 24 hpf, 36 hpf and 48 hpf samples, respectively. Data were normalized using the NormalizeData function and variable features were identified using the FindVariableFeatures function. Cells were then scored based on cell cycle gene expression using the CellCycleScoring function (CC.Difference [S.Score-G2M.Score]), and the data was centered and scaled using the ScaleData function (vars.to.regress = c(“nCount_RNA”, “percent.mt”, “CC.Difference”), features = all.genes). PCA analysis was performed on scaled data and significant principal components were identified by Elbow Plots of the first 100 dimensions. Next, dimensionality reduction was performed using Uniform Manifold Approximation and Projection (UMAP) on the first 40 principal components and clusters were identified using the FindNeighbors (dims = 1:40) and FindClusters functions (24 hpf: res = 1.2; 36 hpf: res = 1; 48 hpf: res = 0.9). Cluster identities were determined by assessing differential expression of known marker genes between different clusters using the cluster biomarkers identified using the FindAllMarkers function (only.pos = T, min.pct = 0.25) (Supplemental Table 1-5).

### Removal of dying cells

After resolution was determined for each data set, the total RNA count, percent mitochondrial gene expression and number of genes detected per cell were re-examined. High mitochondrial gene expression and a low number of feature genes indicative of dying cells were detected in the 24 hpf dataset, cluster 7 and the 36 hpf dataset, cluster 13. These clusters were removed from each dataset leaving 1,826 cells in the 24 hpf data and 2,044 cells in the 36 hpf data set.

### Unsupervised Seurat analysis of integrated data

The three final Seurat objects generated for each stage were integrated into a larger Seurat object through anchor identification using the FindIntergationAnchors and IntergrateData Seurat functions (dims = 1:40) (6264 cells). After integrating the data sets the same downstream analysis was conducted as described in the individual datasets (res = 1.2). Cluster identities were determined by assessing differential expression of known marker genes between different clusters using the cluster biomarkers identified using the FindAllMarkers function (only.pos = T, min.pct = 0.25) (Supplemental Table 4) and comparing previously determined cell identify at each stage.

### Seurat Sub-Clustering MEP Glia and CMNs

Within the integrated data set, cells from the MEP glia and Cranial motor neuronal (CMN) lineages were clustered together. To resolve these populations, the cluster containing these cells was subsetted and sub-clustered by using the FindNeighbors (dims = 1:40) and FindClusters (res = 0.5) Seurat functions. Examination of genes characteristic of MEP glia and CMNs instructed the split in the original cluster, which was then added to the final integrated Seurat object (Fig. S5).

### Seurat Sub-Clustering oligodendrocyte lineage

Within the integrated dataset we noted that some interneuron-like cells and Pre-OPC cells were clustered together. To resolve these populations, the oligodendrocyte lineage clusters (Pre-OL, OPC and Pre-OPC) were subsetted and sub-clustered by using the FindNeighbors (dims = 1:40) and FindClusters (res = 0.5) Seurat functions. Examination of genes characteristic of interneurons and Pre-OPCs instructed the split in the original cluster. The interneuron-like cells were removed from the oligodendrocyte lineage subset (Fig. S8) and trajectory analysis was performed on the adjusted oligodendrocyte lineage.

### Trajectory analysis

#### RNA Velocity

We performed RNA Velocity analysis to infer potential developmental trajectories within the neuronal and oligodendrocyte lineages identified in our integrated and 24 hpf datasets. We used the Python implementation of velocyto (v0.17.13) (La Manno et al., 2018), using the basic ‘run’ subcommand with our CellRanger output and GRCz11 genome assembly. We also opted to mask expressed repetitive elements using the GRCz11 expressed repeat annotation file downloaded from the UCSC genome browser. We analyzed the output loom file using scVelo (v0.2.2) (Bergen et al., 2020), and merged the loom file with a loom file generated with the Seurat as.loom function using our integrated and 24 hpf Seurat objects. The resulting abundance of spliced-to-unspliced RNA was 0.84:0.16 (integrated) and 0.9:0.1 (24 hpf). We used default arguments in order to compute moments for velocity estimation.

#### Slingshot

We also used Slingshot (v1.7.3) (Street et al., 2018) as a second, independent method of trajectory analysis on subsetted neuronal and oligodendrocyte lineages identified in our integrated data set. UMAP reduction was used to determine dimensionality and unbiased lineages were constructed by specifying only a start cluster (pMN1 for the neuronal lineage; Pre-OPC1 for the oligodendrocyte lineage). Lineages and gene expression were visualized using a combination of Slingshot visualization tools, the ggplot2 R package (v3.3.2) and the scater R package (v1.17.4) (McCarthy et al., 2017).

#### Trajectory-based differential expression analysis

To infer differential gene expression in the lineages predicted by our Slingshot analysis, we used a generalized additive model (GAM) with a loess term for pseudotime (gam R package, v1.20) to analyze the top 100 most variable genes across pseudotime. We used a similar testing framework to that illustrated in the online Slingshot tutorial (Street et al., 2018). Heatmaps of the top 100 most variable genes were generated using the heatmap function in base R.

## Acknowledgements

We thank Christina Kearns for isolating cells for scRNA-seq and members of the Appel lab and the Section of Developmental Biology for discussions and advice. Cell sorting was performed by the University of Colorado Cancer Center Flow Cytometry Shared Resource, supported by the Cancer Center Support Grant (P30CA046934). scRNA-seq was performed by the University of Colorado Anschutz Medical Campus Genomics Shared Resource Core Facility, supported by the Cancer Center Support Grant (P30CA046934). Single-cell RNA sequencing and bioinformatics analysis was supported by a pilot award from the University of Colorado RNA Bioscience Initiative. The University of Colorado Anschutz Medical Campus Zebrafish Core Facility was supported by National Institutes of Health grant P30 NS048154. The University of Colorado RNA Bioscience Initiative, the University of Colorado Section of Developmental Biology and the Gates Frontiers Fund provided postdoctoral training support for C.C.W.

## Data availability

The single-cell RNA sequencing data have been deposited in GEO under accession number GSE173350.

R coding scripts are available upon request

## SUPPLEMENTAL DATA

**Fig S1.**
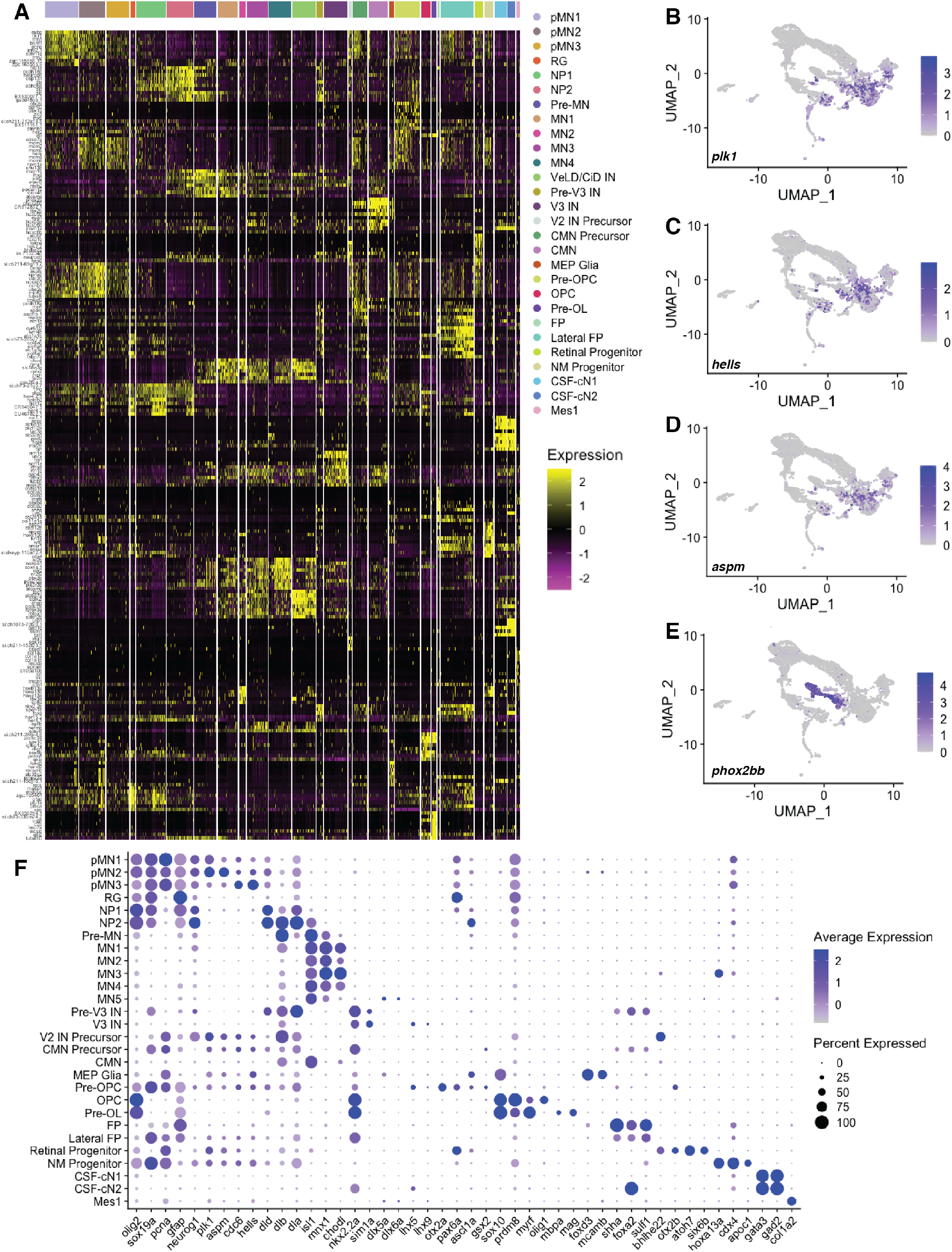
Integrated data cell composition. (A) Heatmap of the top 10 genes significantly enriched in each cluster. Each column represents expression in one cell. (B-E) UMAP feature plots of selected transcripts. Cells are colored by expression level (gray is low, purple is high). Differential expression of cell cycle genes *plk1, hells* and *aspm* distinguish pMN1-3 progenitor populations (B-C). *phox2bb* denotes cranial neuronal precursors and CMN (E). (F) Dot plot identifying molecular markers of each cluster. Dot size portrays the percentage of cells expressing the feature in each cluster. Dots are colored by average expression level (gray is low, purple is high).

**Fig S2.**
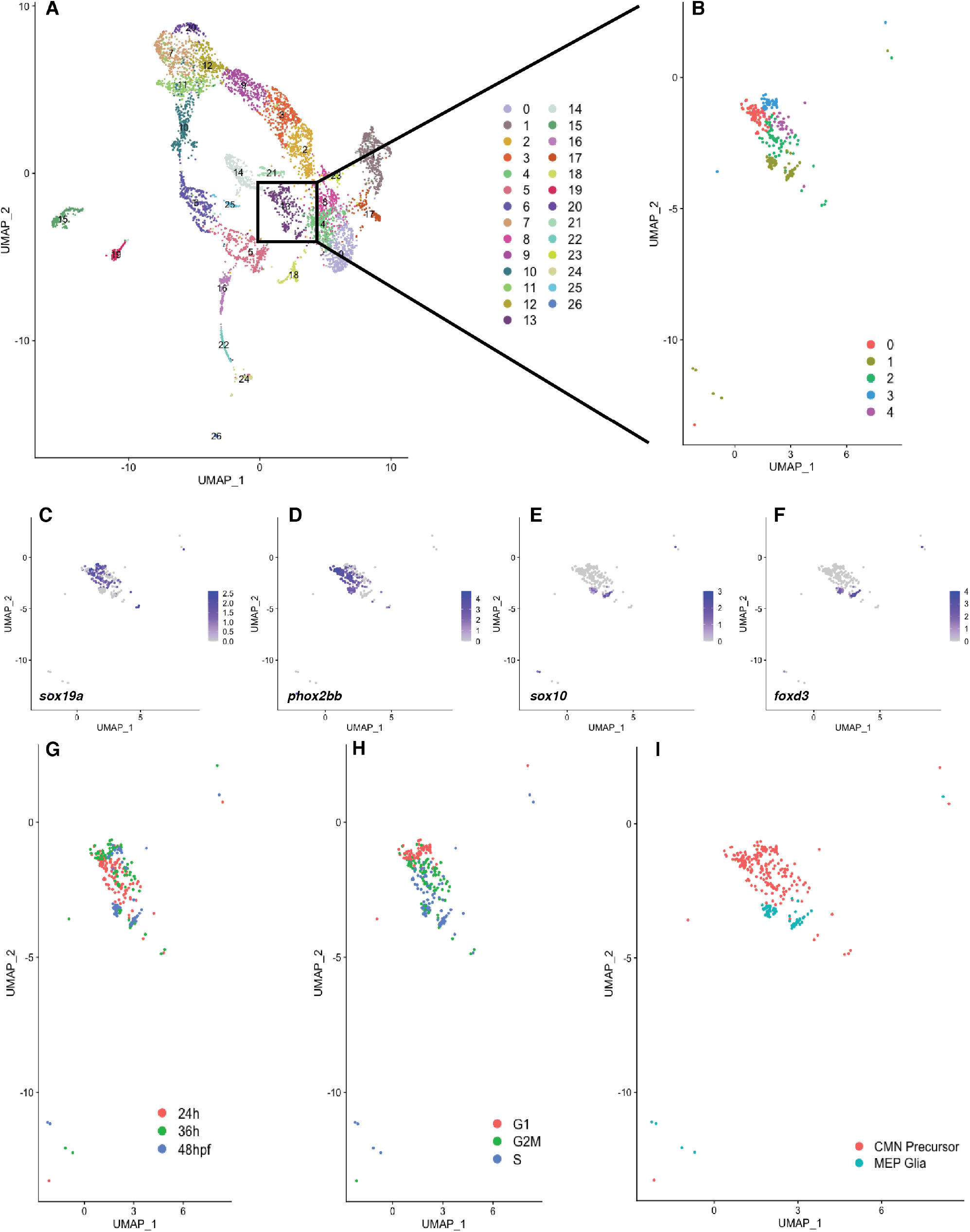
Subcluster analysis reveals CMN Precursors and MEP glia within the integrated data set. (A) UMAP visualization of the scRNA-seq data from *olig2*:EGFP^+^ spinal cord cells obtained from 24, 36 and 48 hpf *Tg(olig2:EGFP)* embryos. Each point represents one cell (n=274). (B) Sub-clustering of cluster 13 resulted in 5 groupings. (C-F) UMAP feature plots of selected transcripts. Cells are colored by expression level (gray is low, purple is high). Co-expression of *sox19a* and *phox2bb* represent CMN precursors (C,D). Co-expression of *sox10* and *foxd3* denote MEP glia (E,F). (G,H) UMAP visualization of cluster 13 sub-clustering. Colors represent sample time points (G) or cell cycle stage (H). (I) UMAP visualization of final sub-clustering determined by differences in expression of select transcripts shown in C-F.

**Fig S3.**
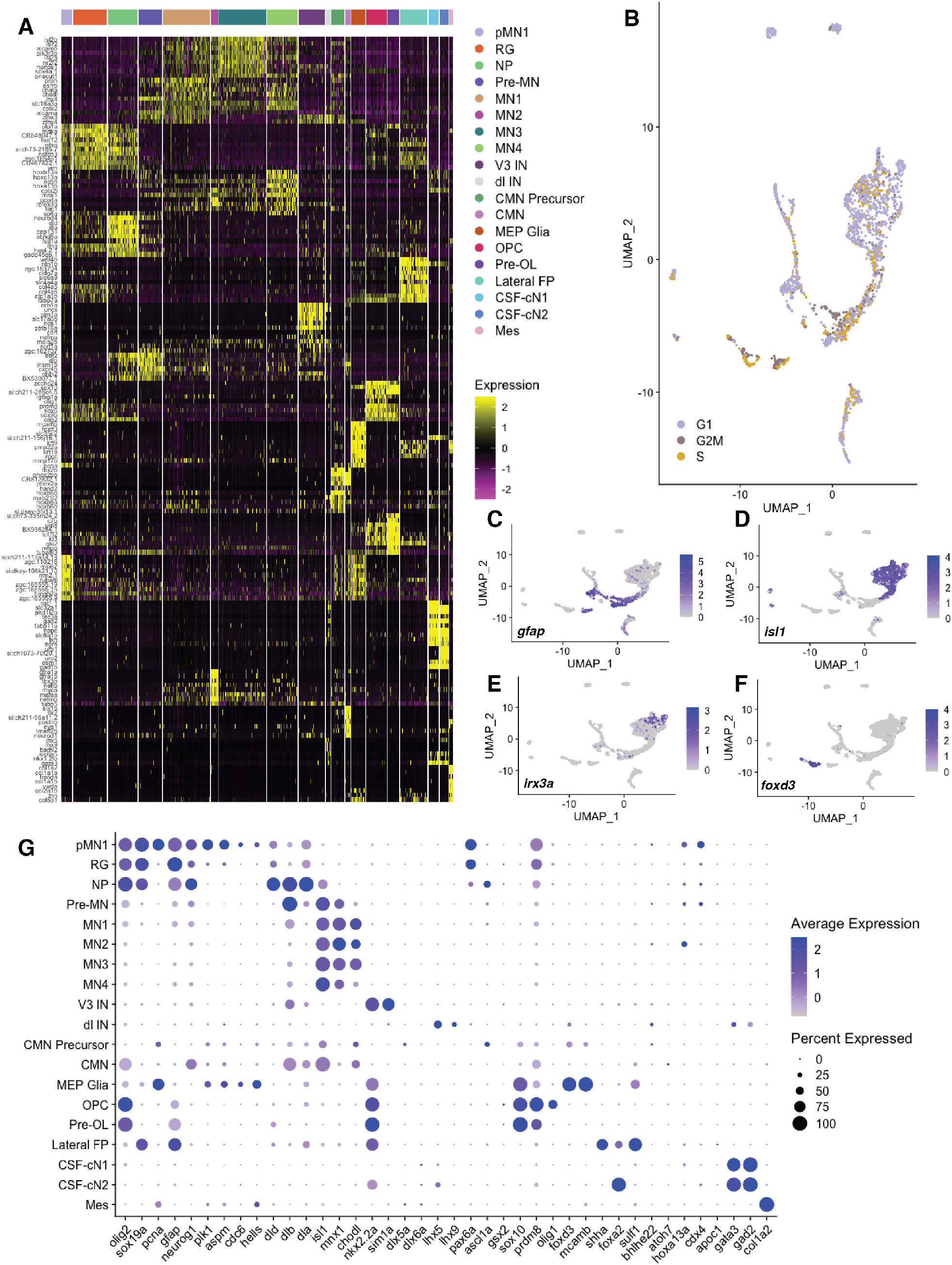
48 hpf cell type composition. (A) Heatmap of the top 10 genes significantly enriched in each cluster of the 48 hpf dataset. Each column represents expression in one cell. (B) UMAP visualization of the scRNA-seq data from *olig2:*EGFP^*+*^ spinal cord cells obtained from 48 hpf *Tg(olig2:EGFP)* embryos. Each point represents one cell (n=2394). Colors represent cell cycle stage. (C-F) UMAP feature plots of selected transcripts. Cells are colored by expression level (gray is low, purple is high). *gfap* marks RG and pMN1 cells (C). *isl1* denotes motor neurons (D) and *irx3a* expression signifies VeLD/CiD INs (E). Expression of *foxd3* illustrates MEP glia (F). (G) Dot plot depicting molecular markers of each cluster. Dot size represents the percentage of cells expressing the feature in each cluster. Dots are colored by average expression level (gray is low, purple is high).

**Fig S4.**
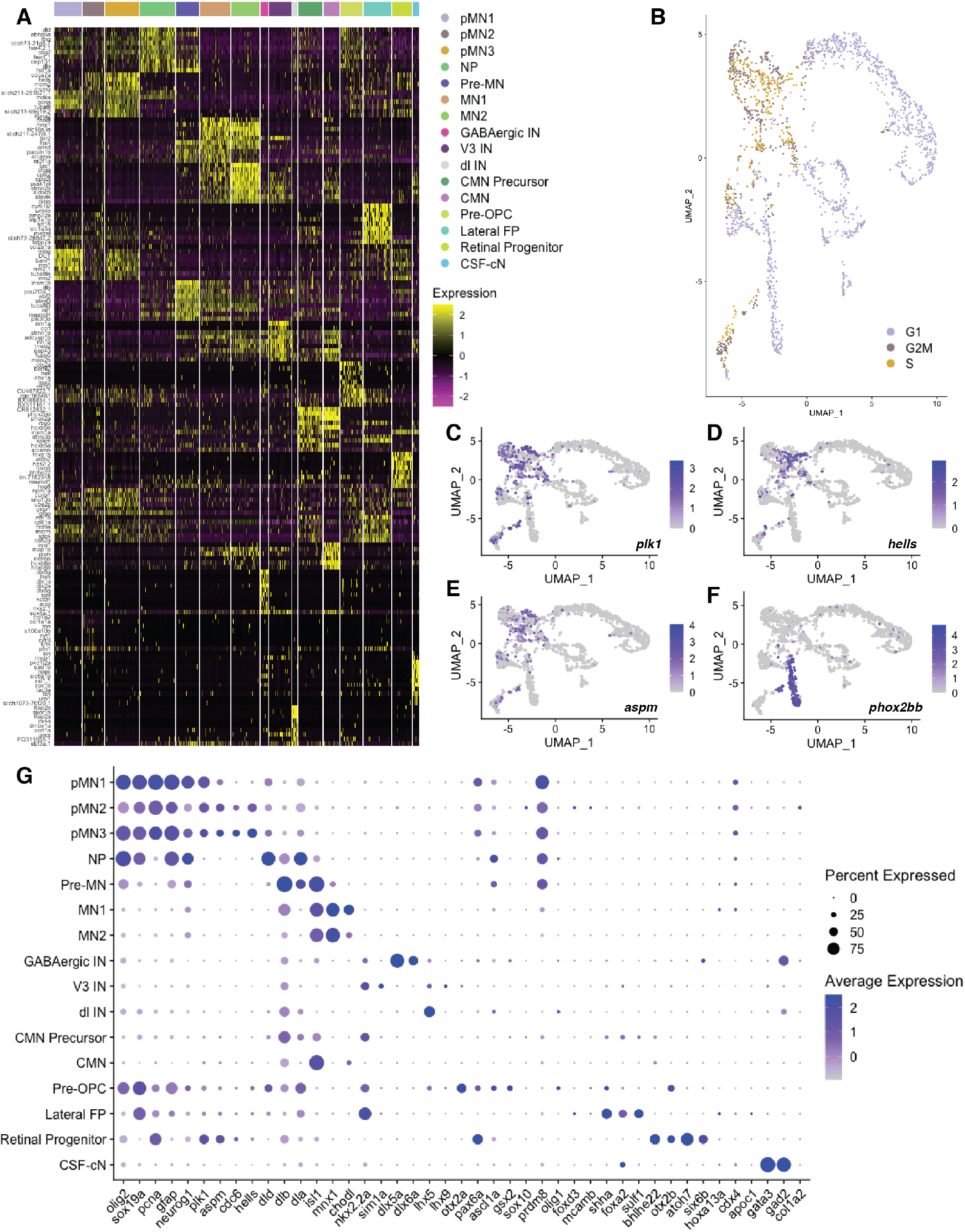
36 hpf cell type composition. (A) Heatmap of the top 10 genes significantly enriched in each cluster of the 36 hpf dataset. Each column represents expression in one cell. (B) UMAP visualization of the scRNA-seq data from *olig2:*EGFP^*+*^ spinal cord cells obtained from 36 hpf *Tg(olig2:EGFP)* embryos. Each point represents one cell (n=2044). Colors represent cell cycle stage. (C-F) UMAP feature plots of selected transcripts. Cells are colored by expression level (gray is low, purple is high). Differential expression of cell cycle genes *plk1, hells* and *aspm* distinguish pMN1-3 progenitor populations (C-E). *phox2bb* denotes cranial neuronal precursors and CMN (F). (G) Dot plot depicting molecular markers of each cluster. Dot size shows the percentage of cells expressing the feature in each cluster. Dots are colored by average expression level (gray is low, purple is high).

**Fig S5.**
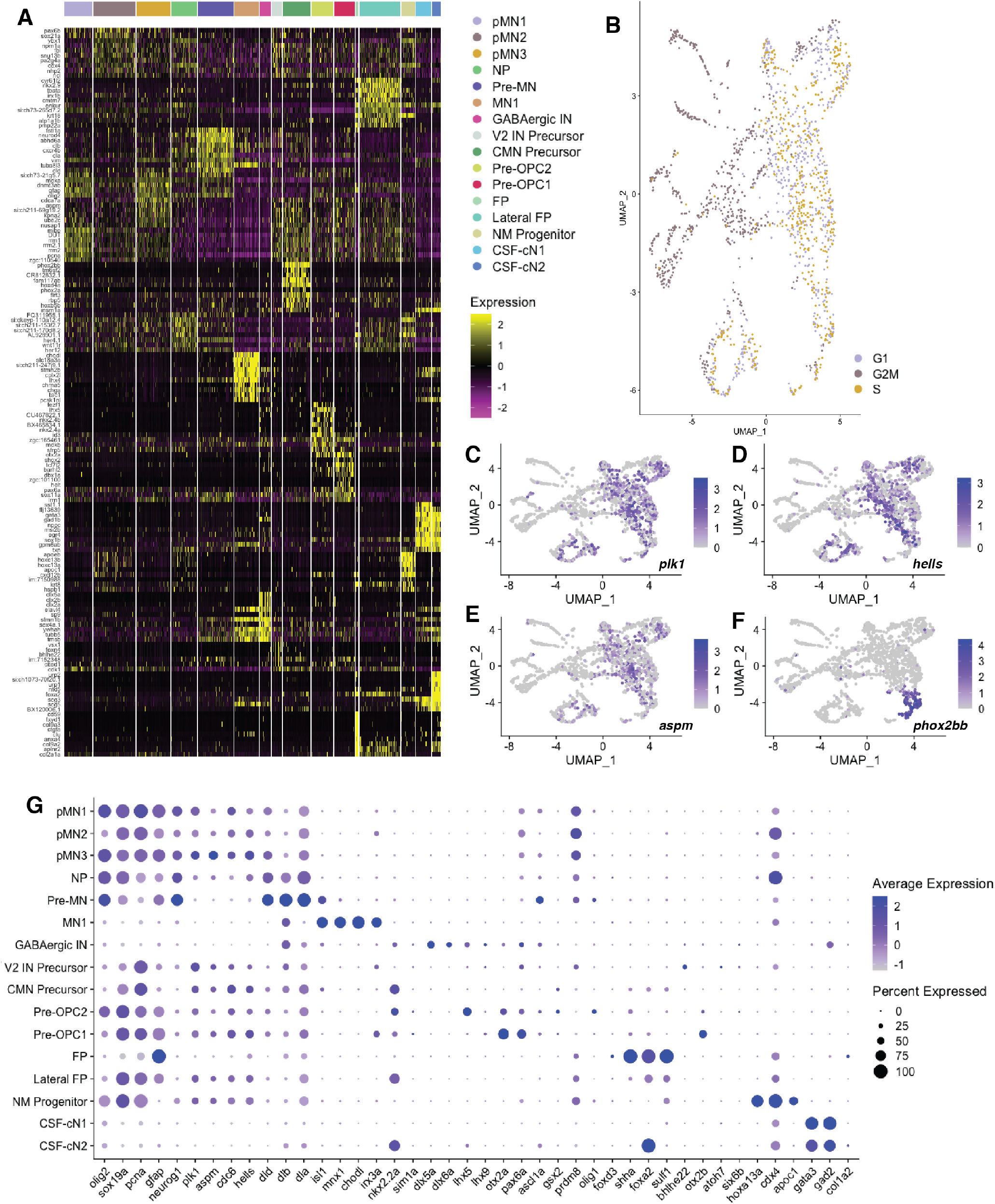
24 hpf cell type composition. (A) Heatmap of the top 10 genes significantly enriched in each cluster of the 24 hpf dataset. Each column represents expression in one cell. (B) UMAP visualization of the scRNA-seq data from *olig2:*EGFP^*+*^ spinal cord cells obtained from 24 hpf *Tg(olig2:EGFP)* embryos. Each point represents one cell (n=1826). Colors represent cell cycle stage. (C-F) UMAP feature plots of selected transcripts. Cells are colored by expression level (gray is low, purple is high). Differential expression of cell cycle genes *plk1, hells* and *aspm* distinguish pMN1-3 progenitor populations (C-E). *phox2bb* denotes cranial neuronal precursors and CMN (F). (G) Dot plot identifying molecular markers of each cluster. Dot size depicts the percentage of cells expressing the feature in each cluster. Dots are colored by average expression level (gray is low, purple is high).

**Fig S6.**
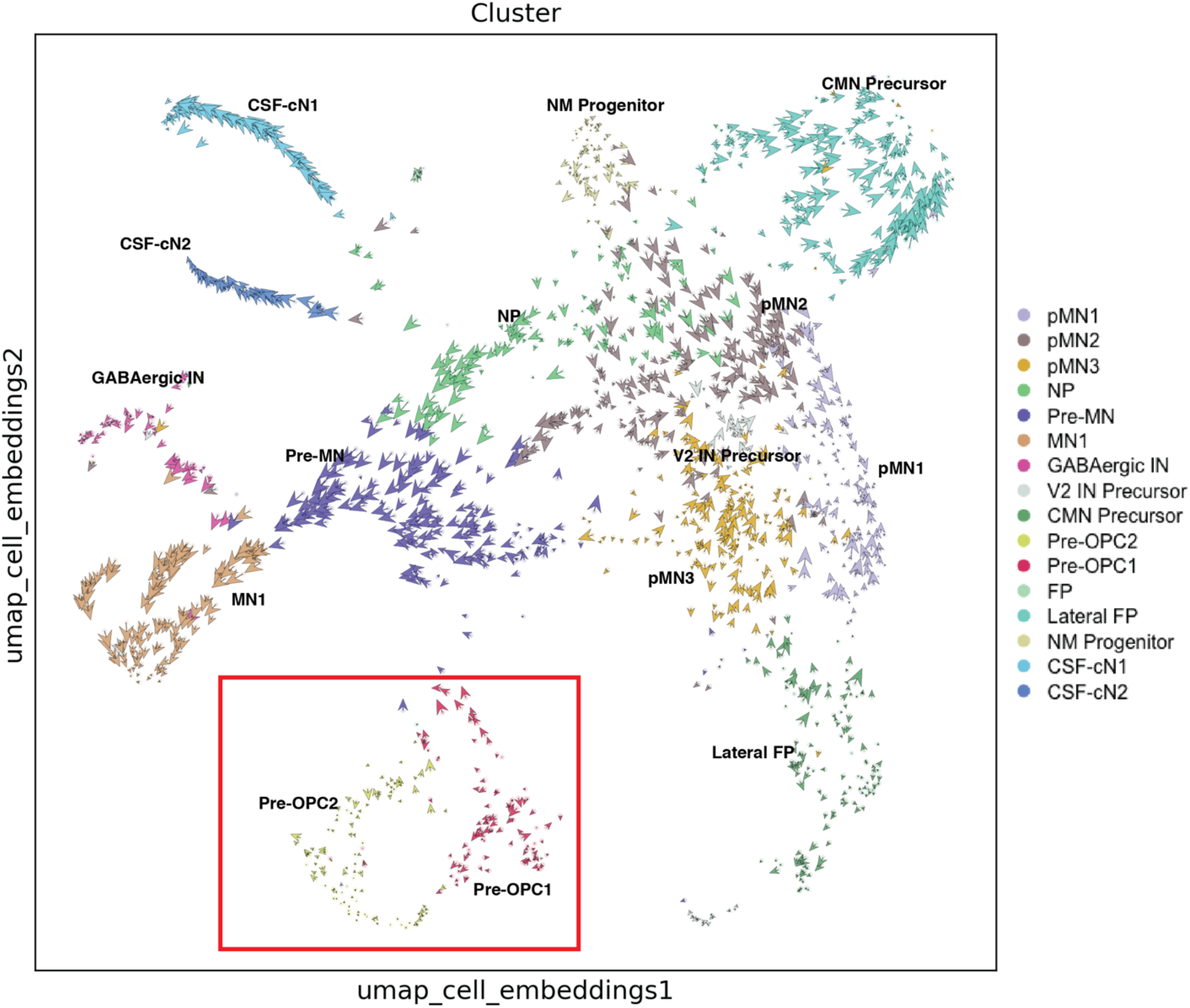
Pre-OPC1 cells precede Pre-OPC2 cells developmentally. Fine-grained resolution of the velocity vector field derived using the scVelo stochastic model, shown at single-cell level. Red box highlights the predicted developmental flow of Pre-OPC1 to Pre-OPC2 cells. Each vector represents the direction and speed of movement of an individual cell.

**Fig S7.**
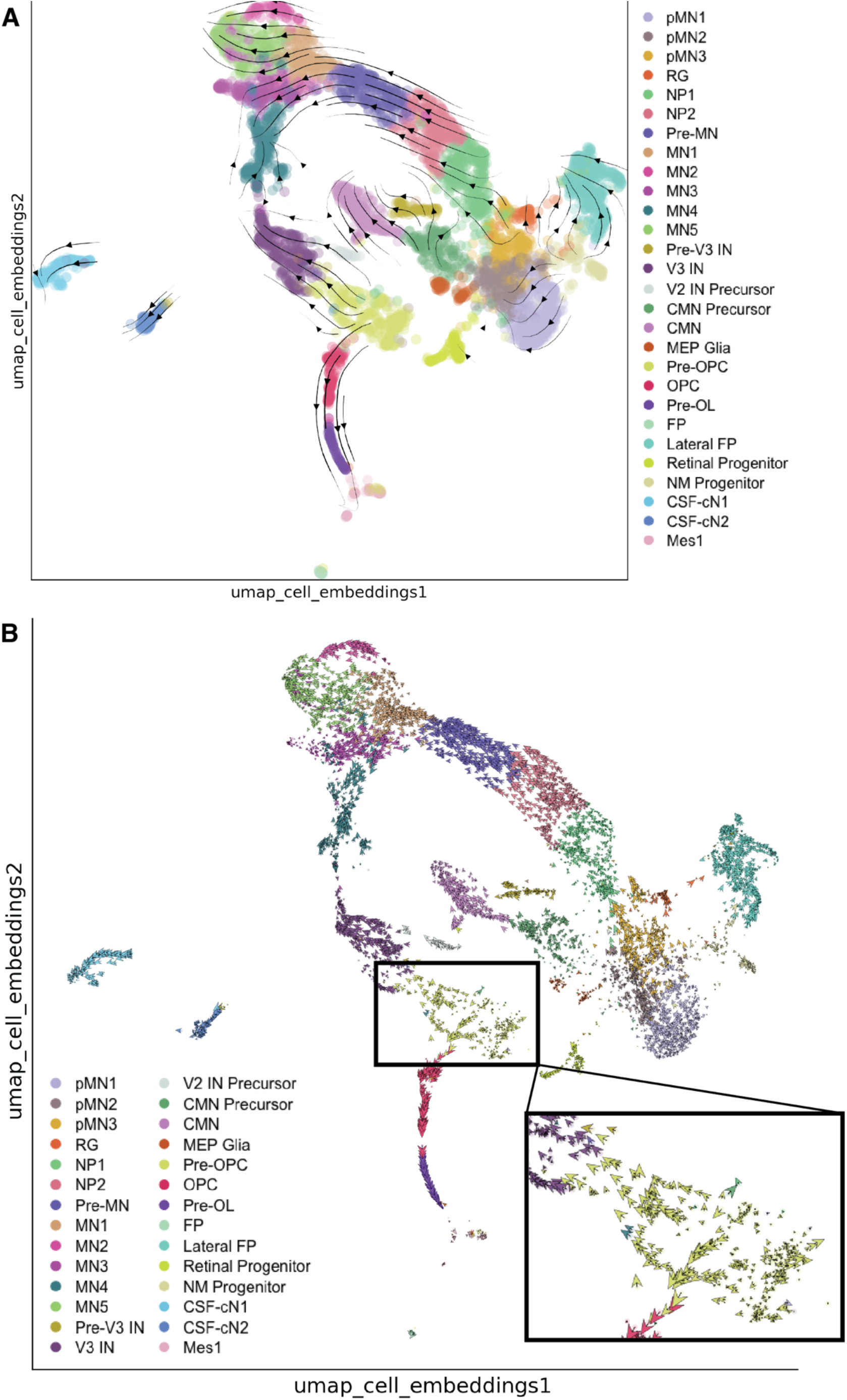
Integrated Pre-OPC cluster includes two diverging trajectories. Velocities derived using the scVelo stochastic model projected onto the UMAP. (A) The main gene-averaged flow visualized by velocity streamlines shows a split within the Pre-OPC cluster. (B) Fine-grained resolution of the velocity vector field in (A) shown at single-cell level. Box highlights diverging trajectories of cells within the Pre-OPC cluster, with one trajectory flowing into the IN lineage and the other flowing into the OLC lineage. Each vector represents the direction and speed of movement of an individual cell.

**Fig S8.**
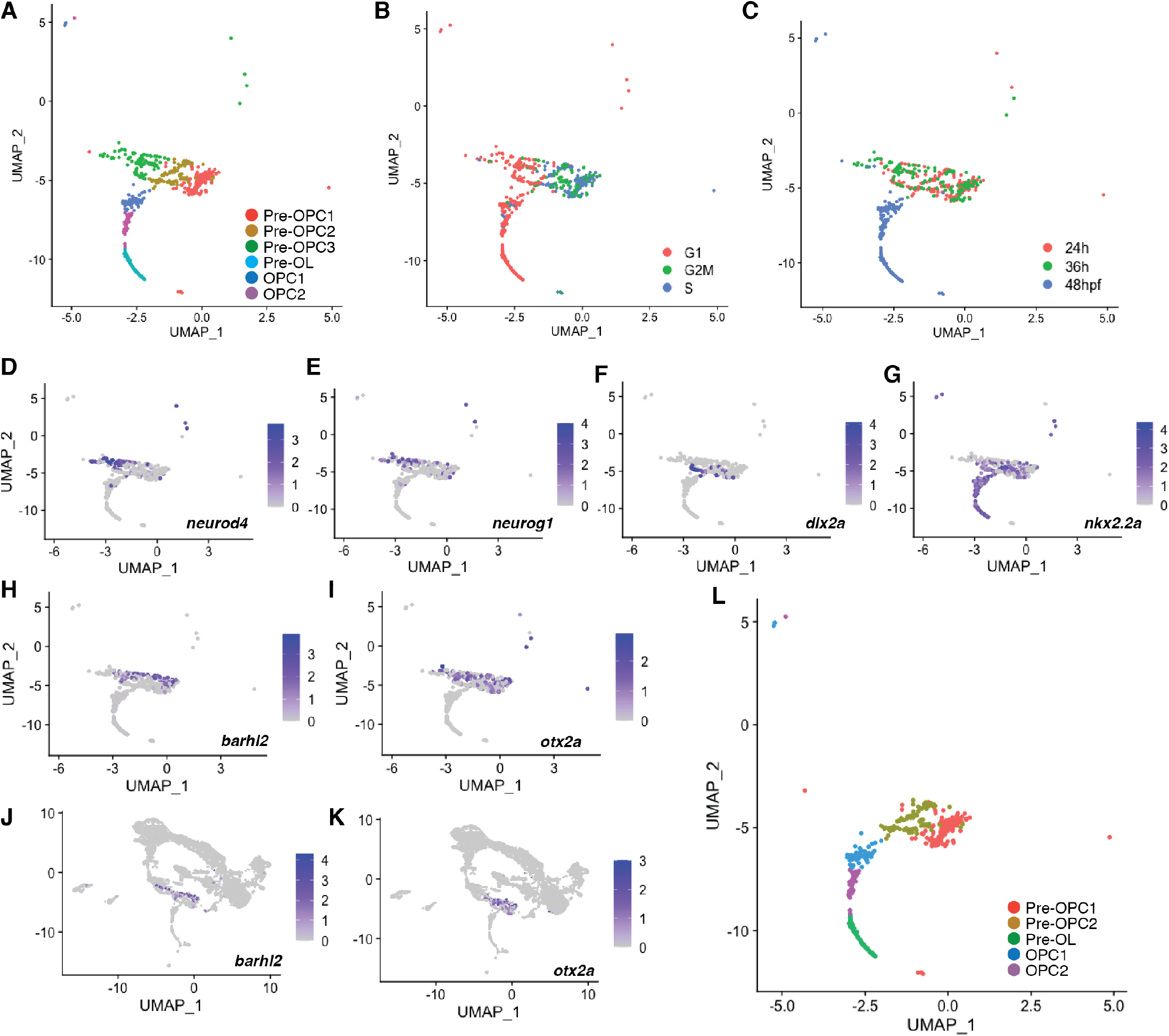
Sub-cluster analysis reveals that Pre-OPCs and IN precursors are grouped in the integrated data set. (A) Sub-clustering of the OLC lineage (n=553) results in 6 clusters. (B-C) UMAP visualization of sub-clustering. Colors represent sample cell cycle stage (B) and time point (C). (D-K) UMAP feature plots of selected transcripts. Cells are colored by expression level (gray is low, purple is high). Combinatorial expression of *neurod4, neurog1* and *dlx2a* denote IN precursors (D-F) and increased expression of *nkx2*.*2a* in Pre-OPC1 and Pre-OPC2 sub-clusters mark the oligodendrocyte lineage cells. (G). *barhl2* and *otx2a* are uniquely expressed in the Pre-OPC1-3 (H,I) lineages within the integrated dataset (J,K). (L) UMAP visualization of final sub-clustering determined by differences in expression of select transcripts shown in D-G.

**Fig S9.**
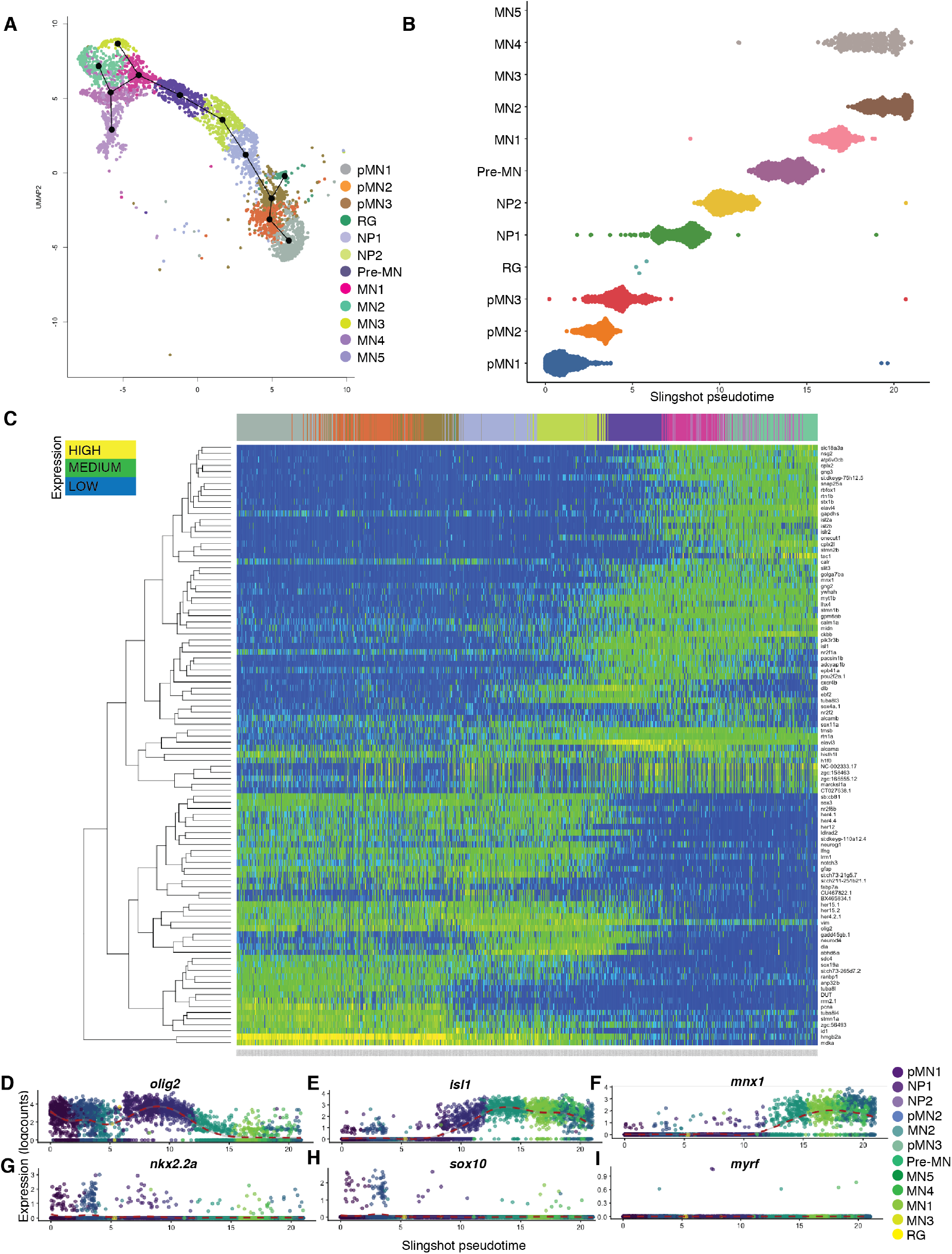
Pseudotime analysis of neuronal lineage pMN cells. (A) UMAP of sub-clustered neuronal lineage (n=3632) from integrated data set depicting the lineage trajectory predicted by Slingshot analysis. (B) Depiction of cells within each sub-cluster from the Slingshot-predicted lineage along pseudo-temporal ordering. Each dot represents a cell and its predicted temporal position in development. (C) Heatmap showing top 100 differentially expressed transcripts based on the Slingshot-predicted oligodendrocyte lineage (left = pMN1 Progenitors, right = MNs). Dendrogram displays hierarchical clustering; each column represents the relative expression of each labeled transcript in a single cell. (D-I) Scater smooth expression plots for transcripts related to neuronal (D-F) or oligodendrocyte (D, G-I) development within each subcluster along Slingshot pseudo-temporal ordering. Each point represents the expression of each transcript within a single cell. *olig2* is expressed in pMN1-3 and NPs but is downregulated in mature MNs (D). *isl1* is expressed by NP1/2 (E). Mature MNs maintain expression of *isl1* and upregulate *mnx1* (E,F). A small number of pMN progenitors express *sox10* and *nkx2*.*2a* whereas most NPs and MNs do not express these transcripts (G-H). Most neuronal lineage cells do not express *myrf* (I).

## KEY RESCOURCES TABLE

Developmental Biology Formatted

KEY RESOURCES TABLE: Developmental Biology Formatted
| Reagent or resource | Source | Identifier |
| --- | --- | --- |
| Chemicals, Peptides, and Recombinant Proteins |  |  |
| Molecular Grade Water | Corning Cellgro | Cat# 46-000-CV |
| Sucrose | BDH | Cat# 57-50-1 |
| Agar | Fisher BioReagents | BP1423-500 |
| Tricaine | Western Chemicals Inc | TRICMGR0100 |
| Paraformaldehyde | EMS | Cat# 15714 |
| Agarose | Fisher BioReagents | Cat# BP160-500 |
| Vectashield | Vector Laboratories | Cat# H-1000 |
| NaCl | Omnipur | Cat# 4099-06 |
| KCl | Calbio Chem | Cat# 7300 |
| CaCl<sub>2</sub> | EMD | CS0130-2 |
| MgSO<sub>4</sub> | EMD | MX0070-1 |
| BI Protease | Sigma | P5380 |
| Critical Commercial Assays |  |  |
| RNAscope® Multiplex Fluorescent Reagent Kit V2 | ACD Bio | 323100 |
| Opal 7 Kit | PerkinElmer | NEL797001KT |
| Deposited Data |  |  |
| Experimental Models: Organisms/Strains |  |  |
| Zebrafish: *Tg(olig2:EGFP)<sup>vu12</sup>* | (Shin et al., 2003) | ZFIN ID: ZDB-ALT-041129-8 |
| Zebrafish: AB |  | ZFIN ID: ZDB-GENO-960809-7 |
| Oligonucleotides |  |  |
| RNAscope Probe-Dr-olig2 | ACD | 559211 |
| RNAscope Probe-Dr-ascl1a-C1 | ACD | 583661 |
| RNAscope Probe-Dr-neurog1-C2 | ACD | 505081-C2 |
| RNAscope Probe-Dr-gsx2-C3 | ACD | CUSTOM PROBE |
| Software and Algorithms |  |  |
| ImageJ Fiji software | NIH | https://imagej.nih.gov/ij/download.html |
| ZEN Black | Carl Zeiss | https://www.zeiss.com/microscopy/us/products/microscope-software.html |
| Zen Blue | Carl Zeiss | https://www.zeiss.com/microscopy/us/products/microscope-software.html |
| R Studio (4.0.1) | R Studio | https://rstudio.com/ |
| Seurat (3.2.1) | Satija Lab | https://satijalab.org/seurat/ |
| Slingshot (1.7.3) | (Street et al., 2018) | https://bioconductor.org/packages/release/bioc/html/slingshot.html |
| scater (1.17.4) | (McCarthy et al., 2017) | http://bioconductor.org/packages/release/bioc/html/scater.html |
| MAST |  | https://doi.org/10.1186/s13059-015-0844-5 |
| Other |  |  |

